# Dirus Complex Species Identification PCR (DiCSIP) improves identification of *Anopheles dirus* complex from Greater Mekong Subregion

**DOI:** 10.1101/2023.10.09.561471

**Authors:** Manop Saeung, Jutharat Pengon, Chatpong Pethrak, Sarunya Thaiudomsap, Suthat Lhaosudto, Atiporn Saeung, Sylvie Manguin, Theeraphap Chareonviriyaphap, Natapong Jupatanakul

## Abstract

**Background:** The *Anopheles dirus* complex plays a significant role as malaria vectors in the Greater Mekong Subregion (GMS), with varying degrees of vector competence among species. Accurate identification of sibling species in this complex is essential for understanding malaria transmission dynamics and deployment of effective vector control measures. This becomes increasingly crucial as the GMS advances towards malaria elimination while facing with the emergence of zoonotic simian malaria transmission. However, the original molecular identification assay, Dirus Allele Specific PCR (AS-PCR) targeting the ITS2 region have pronounced non-specific amplifications leading to ambiguous results and misidentification of the sibling species. This study investigates the underlying causes of these inconsistencies and develops new primers for accurate identification of species within the *Anopheles dirus* complex.

**Methodology/Principal findings:** Despite several optimizations by reducing primer concentration, decreasing thermal cycling time, and increasing annealing temperature, the Dirus AS-PCR continued to produce inaccurate identifications, particularly for *Anopheles dirus*, *Anopheles scanloni*, and *Anopheles nemophilous*. Subsequently, *in silico* analyses pinpointed problematic primers with high GC content and multiple off-target binding sites. Through a series of *in silico* analyses and laboratory validation, a new set of primers for Dirus Complex Species Identification PCR (DiCSIP) has been developed. DiCSIP primers improve specificity, operational range, and sensitivity for accurate identification of five complex member species found in the GMS. Validation with laboratory and field *An. dirus* complex specimens demonstrated that DiCSIP could correctly identify all samples while the original Dirus AS-PCR misidentify *An. dirus* as other species when used with different thermocyclers.

**Conclusion/Significance:** The DiCSIP assay offers a significant improvement in *An. dirus* complex identification, addressing challenges in specificity and efficiency of the previous ITS2-based assay. This new primer set provides a valuable tool for accurate entomological surveys, supporting effective vector control strategies to reduce transmission, and prevent the re-introduction of malaria in the GMS.

**Author Summary:** Several species of *Anopheles* mosquitoes belong to species complexes due to their indistinguishable morphology from closely related sibling species. However, members of the same species complex may exhibit varied vectorial capability, i.e. the ability to spread human pathogens, in this case malaria parasites, ranging from dominant vector to non-vector. The Dirus AS-PCR molecular assay to identify species within the *An. dirus* complex, a significant malaria vector species in the GMS, can produce inconsistent PCR results leading to misidentification. Furthermore, our analysis of *An. dirus* ITS2 sequences in the NCBI database indicated misidentification between these sibling species suggesting the need for a new set of primers to improve reproducibility and sensitivity in identifying members of the *An. dirus* complex. This study presents thorough analyses of the existing primers that cause difficulties in correct amplification and a development of a new set of primers for Dirus Complex Species Identification PCR (DiCSIP) assay. The DiCSIP assay offers several advantages over the original Dirus AS-PCR. It provides a higher specificity, sensitivity and wider operational range allowing for the use of the DNAzol direct reagent to process mosquito samples without the need for DNA extraction, saving both time and cost in sample processing. Our study provides a valuable molecular tool for entomological surveys, which is crucial for effective vector control measures in the GMS.

## 1. Introduction

In recent decades, countries in the Greater Mekong Subregion (GMS), particularly Thailand, have made significant progress in reducing the number of endemic malaria cases (1). With declining malaria cases, the focus has shifted towards the goal of malaria elimination. At this stage, entomological data becomes increasingly crucial to evaluate areas with potential malaria transmission and to prevent disease recurrence, especially considering the threat of zoonotic transmission of simian malaria spill over to humans (2).

Among more than 530 species of *Anopheles* mosquitoes, only tens of them are vectors of pathogens responsible for malaria and other vector-borne diseases (3, 4). Within this genus, certain species belong to a species complex due to the indistinguishable morphological characters of closely related sibling species, while their vectorial capacity ranges from main vectors to non-vectors. Therefore, identifying species among *Anopheles* mosquitoes to evaluate transmission risks represents a real challenge. Prior to advancements in molecular genetics, there were indications of misidentification at the species complex level based solely on morphological characteristics (5–8). Subsequently, genetic information-based techniques have been employed to complement morphological identification. Polymerase chain reaction (PCR) assays has been developed and utilized for identifying mosquitoes within *Anopheles* complexes such as *Anopheles dirus*, *Anopheles minimus*, *Anopheles barbirostris*, *Anopheles leucosphyrus* (9–12). The internal transcribed spacer 2 (ITS2) is widely utilized as the primary DNA region for these purposes, mainly because it contains conserved regions suitable for designing universal primers and offers sufficient variability to distinguish even closely related species (13). Another advantage of the ITS2 gene over other commonly used molecular markers is the ability to distinguish species based on different amplicon sizes, thus eliminating the need for costly sequencing.

The *Anopheles dirus* complex has been considered a medically important taxa in the GMS due to its ability to transmit *Plasmodium* parasites, pathogens responsible for human malaria (1, 14, 15). Furthermore, the Dirus complex has been reported to be a vector of simian malaria, *P. knowlesi*, which currently causing a concern of zoonotic transmission in the GMS countries (1, 16, 17). The taxonomic hierarchy of the Dirus complex was recognized as belonging to Subgenus *Cellia*, Neomyzomyia Series, Leucosphyrus Group, Leucosphyrus Subgroup, (18). The distribution range of the Dirus complex is mainly in Southeast Asia, including Thailand (19). The complex comprises eight sibling species, six of which can be found in the GMS, including *An. dirus* (former *An. dirus* A), *Anopheles cracens* (former *An. dirus* B), *Anopheles scanloni* (former *An. dirus* C), *Anopheles baimaii* (former *An. dirus* D), *Anopheles nemophilous* (former *An. dirus* F), and *Anopheles* aff. *takasagoensis* (18). For the other two sibling species, *An. elegans* is present in southern India, while *An. takasagoensis* has only been found in Taiwan (4, 20). In Thailand, five species are present as listed above, except *Anopheles* aff. *takasagoensis,* which is restricted to northern Vietnam (21). Among all eight species, two are recognized as major malaria vectors, namely *An. dirus* and *An. baimaii* (22–25), while the others have been described as secondary or non-vector. Due to overlapping larval habitats and sympatric occurrence of the five sibling species found in Thailand, relying solely on ecological characteristics and/or morphological characters becomes insufficient to distinguish them. Therefore, molecular-based techniques are essential for precise identification of primary malaria vectors and for categorizing risk areas based on the geographical distribution of these vectors (22).

The *An. dirus* complex Allele-Specific multiplex PCR (Dirus AS-PCR), targeting the ITS2 gene, was designed for differentiating five sibling species of the Dirus complex in the GMS. The assay uses one universal forward primer (D-U) and four species-specific reverse primers, namely D-AC (specific to *An. dirus* and *An. scanloni*), D-B (*An. cracens*), D-D (*An. baimaii*), and D-F (*An. nemophilous*). Since its development in 1999, this technique, has been adopted for entomological surveys in this region (9–12). However, difficulties in interpretation of PCR results due to non-specific or inefficient amplification have been reported (26).

In this study, we describe efforts to optimize reaction conditions for the Dirus AS-PCR to improve specificity. Additionally, we performed *in silico* analyses, including binding site identification and amplicon predictions, to identify the causes of difficulties in the assay. Furthermore, we developed a new PCR assay called Dirus Complex Species Identification PCR (DiCSIP) to enhance reproducibility and sensitivity of *An. dirus* complex identification.

## 2. Materials and Methods

### 2.1 *Anopheles dirus* complex samples

Laboratory colonies and field-derived mosquitoes belonging to the *An. dirus* complex were used in this study. *An. dirus* samples were obtained from the insectary at the Department of Entomology, Faculty of Agriculture, Kasetsart University (KU). *An. cracens* laboratory samples were obtained from the insectary of the Armed Forces Research Institute of Medical Sciences (AFRIMS) and Chiang Mai University. Field-collected *An. dirus* complex samples were obtained from previous field surveys conducted in Kanchanaburi, Prachinburi, Ranong, and Sisaket Provinces, Thailand. The samples were morphologically identified using a pictorial key under stereomicroscope (23). Due to the lack of field samples of *An. scanloni* and *An. nemophilous*, the gene constructs were synthesized based on their ITS2 sequences obtained from the previous publication describing the original Dirus AS-PCR (9) using Genscript service.

### 2.2 DNA extraction

DNA extraction was conducted using either the Zymo Quick-DNA extraction kit (Zymo research, Cat# D3024) for high purity DNA or DNAzol-direct Reagent (MRC Inc, Cat# DN131) for high-throughput DNA extraction with field samples following the manufacturers’ protocol.

For Zymo Quick-DNA extraction kit, the whole body of each mosquito was homogenized using 0.5 mm sterile glass beads with Bullet Blender homogenizer (NextAdvance) or plastic pestles in 500 µl of genomic lysis buffer. The debris was pelleted by centrifugation at 12,000×g for 5 min, and the supernatant was transferred to a Zymo-Spin III filter in the collection tube and centrifuged at 8,000×g for 1 min. Afterward 1,200 µL Genomic Lysis Buffer was added to the filtrate in the collection tube. Then, the mixture was transferred to a Zymo-Spin IICR column in a collection tube and centrifuged at 10,000×g for 1 min. DNA on the column was washed with 200 µl of DNA Pre-Wash Buffer, followed by washing with 500 µl of g-DNA Wash Buffer. Finally, the column DNA was eluted with 50 µl of DNase/RNase free water. DNA concentration was measured using Qubit dsDNA HS Assay Kit (Invitrogen, Cat# Q32851). The eluted DNA was used immediately or stored at -20°C until further analysis.

DNAzol-direct reagent was used to process laboratory and field samples for Dirus AS-PCR and DiCSIP assay validation. Briefly, two mosquito legs were homogenized in 30 μL of DNAzol-direct reagent using 0.5 mm sterile glass beads with Bullet Blender homogenizer (NextAdvance) or plastic pestles. Tissue debris was pelleted by centrifugation at 8,000xg for 3 min, and the supernatant was used directly in PCR reactions or stored at -20°C until further use.

### 2.3 PCR conditions and optimization

#### 2.3.1 The original Dirus AS-PCR condition

The Dirus AS-PCR was conducted according to a published protocol (9). Briefly, PCR was set up in the total reaction volume of 12.5 μL containing a 2.5 ng of genomic DNA (gDNA) or 0.1 ng of DNA from plasmid harboring ITS2 gene (pDNA), 1X Gotaq Buffer, 1.25 units of Promega GoTaq Flexi DNA Polymerase (Promega, Cat#M8295), 2 mM MgCl_2_, 10% Dimethyl sulfoxide (DMSO), 0.2 mM dNTPs, and 1 μM each of primer D-U, D-AC, D-B, D-D, and D-F. Invitrogen Taq DNA polymerase (Invitrogen, Cat#11615-010) was also used to set up a PCR reaction by replacing the reaction buffer and enzyme while maintaining other reagents. Preparation of PCR reaction was conducted on ice to prevent non-specific amplification during reaction set up. The thermal cycling condition consists of 5 min denaturation step at 94°C, followed by 35 amplification cycles of denaturation at 94°C for 1 min, annealing temperature (Ta) 51°C for 1 min, extension at 72°C for 2 min, and a final extension step at 72°C for 10 min.

PCR products were separated and visualized using 2% agarose gel electrophoresis in 1X Tris-acetate-EDTA (TAE). A volume of 3 µl of each PCR product was mixed with 5X DNA loading dye then individually loaded into each well, and 1.5 µl of a 50 bp DNA ladder (SMOBIO, Cat# DM1100) was used as a marker. The agarose gel was run for 28 min at 100 volts with Mupid-exU electrophoresis system. Then, PCR products on agarose gel were stained with Ethidium bromide for 4 min and visualized under UV light using Syngene G:Box Chemi-XX9-F0.8 imager.

#### 2.3.2 Modifications of the original Dirus AS-PCR

Reducing the primer concentration: the Dirus AS-PCR was conducted as described in 2.3.1 but with final primer concentration reduced from 1 μM to 0.2 μM. Thermal cycling was also shortened with a denaturation at 94°C for 2 min, followed by 35 cycles of denaturation at 94°C for 20 sec, annealing with Ta increased from 51°C to 55°C or 60°C for 20 sec, and extension at 72°C for 1 min, followed by final extension step at 72°C for 5 min. Specific information of the reaction set up and thermal cycling conditions are described in each specific experiment.

### 2.4 Bioinformatic analyses and primer design

Primer-BLAST (27) was used to analyze potential amplicon and species-specificity of the universal forward and species-specific reverse primer pairs against nucleotide (nr) database of the *Anopheles* genus (tax id:7164) in the National Center for Biotechnology Information (NCBI). The parameters for the alignment were set with a product size range of 70 to 1,000 bp and melting temperatures (Tm) at 60 ± 3 °C (***Table 1***).

**Table 1.**
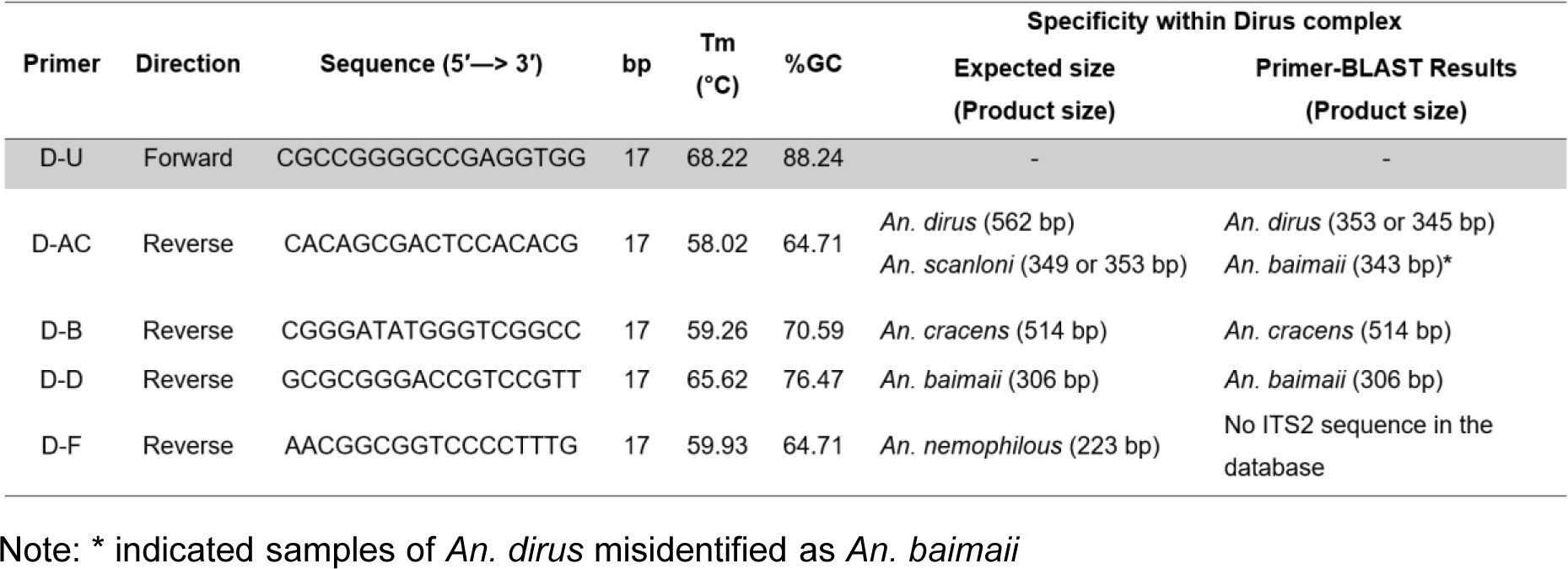
Bioinformatic analyses of Dirus AS-PCR primers. The primer properties were predicted in Benchling online tools and specificity for PCR amplification was analyzed in Primer-BLAST. The search criteria were 70-1,000 bp product size, melting temperatures (Tm) at 60 ± 3 °C, *Anopheles* (taxid:7164).

Benchling online tool (https://www.benchling.com/) was used to analyze primer properties as well as potential binding sites to explain inconsistencies in PCR results. Furthermore, new primers were designed using a primer design tool on Benchling. The DNA sequences of *An. scanloni* and *An. nemophilous* were retrieved from published article (9). DNA sequences of *An. dirus* (Accession number: MW647457), *An. cracens* (Accession number: MG008574), and *An. baimaii* (Accession number: MN152993) were obtained from the GenBank database. Multiple sequence alignments (MSA) were conducted to determine regions appropriate for a new universal forward primer, as well as species specific forward and reverse primers. The parameters for primer design include: 1) nucleotide length with 16-22 bases, 2) G/C content of 57-77%, 3) melting temperature (Tm) of 50-63°C (***Table 2***). A new universal forward primer (DiCSIP-Uni-Fwd) was designed at the conserved region among the five species of the *An. dirus* complex. Whereas new species-specific reverse primers (DiCSIP-Rev) were designed at regions distinct for each species. Due to high similarity between *An. dirus* and *An. scanloni* ITS2 several forward and reverse primers were designed to differentiate these two species (***Supplementary Tables S1***). Potential binding sites of each primer on *An. dirus* complex ITS2 were analyzed on Benchling. Additionally, primer-BLAST was used to identify potential amplicons from new primers as described above (***Table 2***).

**Table 2.**
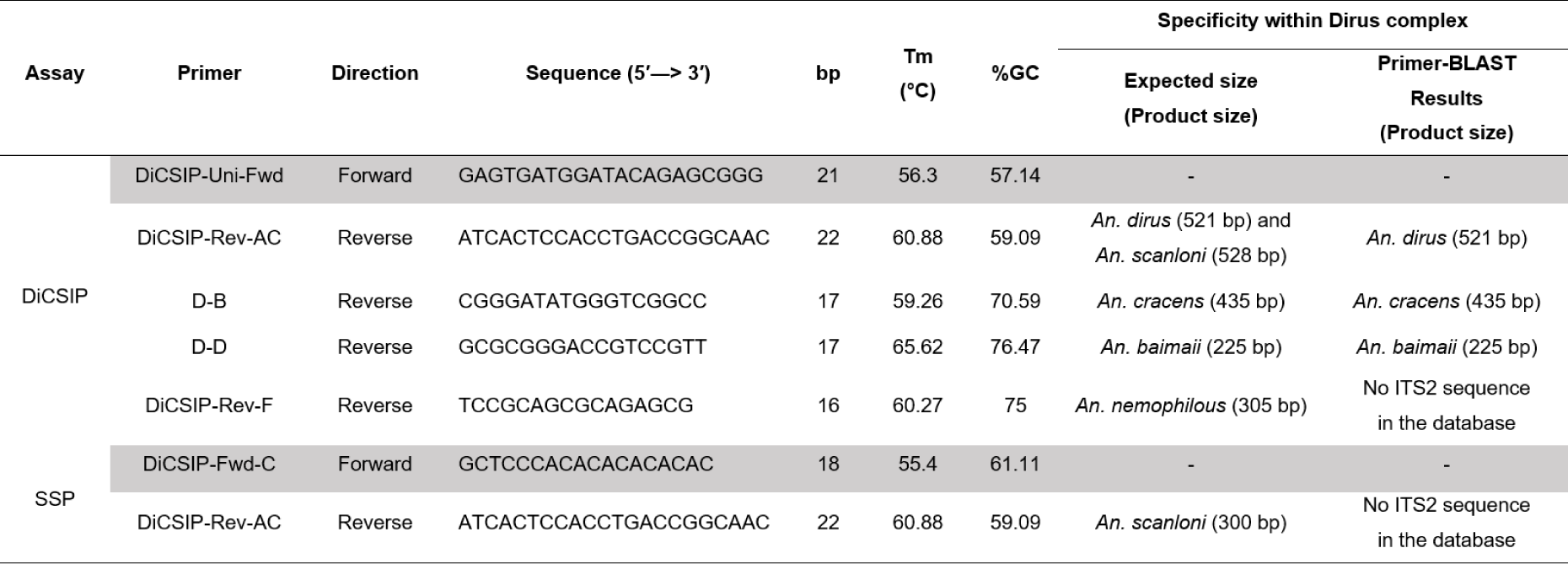
Sequences and bioinformatic analyses of the new DiCSIP primers to distinguish five member species within the *An. dirus* complex. Primer properties were predicted in Benchling online tools and specificity for PCR amplification were analyzed in Primer-BLAST. The search criteria were 70-1,000 bp product size, melting temperatures (Tm) at 60 ± 3 °C, *Anopheles* (taxid:7164), SSP: Scanloni-specific PCR.

### 2.5 Validation of new DiCSIP primers

After *in silico* analyses, new primers were validated for specificity by PCR as followed:

#### 2.5.1 DiCSIP universal forward primer

The DiCSIP-Uni-Fwd primer was validated with four original Dirus AS-PCR reverse primers (9) to confirm whether the replacement of the forward primer improves specificity. The DiCSIP-Uni-Fwd primer was subsequently validated by multiplex PCR with four original Dirus AS-PCR species-specific reverse primers D-AC, D-B, D-D, and D-F. PCR reaction was performed in a volume of 12.5 μL in a final content of 2.5 ng gDNA or 0.1 ng pDNA, 1X Gotaq Buffer, 1.25 units of Promega GoTaq Flexi DNA Polymerase, 2 mM MgCl2, 10% Dimethyl sulfoxide (DMSO), 0.2 mM dNTPs, and 0.2 μM of each primer. The thermal cycling condition consists of 2 min denaturation step at 94°C, followed by 35 amplification cycles of denaturation at 92°C for 20 sec, annealing temperature (Ta) 60°C for 20 sec, extension at 72°C for 1 min, and a final extension step at 72°C for 5 min.

#### 2.5.2 DiCSIP species specific reverse primers

Based on the data from bioinformatic analysis (***Table 1***) and preliminary results, it was found that reverse primers, D-AC and D-F caused difficulties in species identification. After designing the new DiCSIP species-specific reverse primers, they were used in a single-plex PCR reaction together with the DiCSIP-Uni-Fwd primer using the same reaction setup and thermal cycling conditions as described in section 2.5.1, except for a reduction of DMSO final concentration from 10% to 4%.

#### 2.5.3 Scanloni-Specific PCR (SSP)

*An. scanloni*-specific forward primer (DiCSIP-Fwd-C) was validated together with *An. dirus/scanloni*-specific reverse primer (DiCSIP-Rev-AC) using the PCR condition as described in the section 2.5.2 except for a reduction of DMSO from 10% to 4% and an increased Ta of 62°C.

### 2.6 Optimized conditions for DiCSIP assay

The DiCSIP assay consists of two PCR reactions. The first reaction is used to differentiate *An. dirus*/*scanloni*, *An. cracens*, *An. baimaii*, and *An. nemophilous*, based on their respective amplicon sizes. The PCR reactions consist of 1X Gotaq Buffer, 1.25 units of GoTaq® DNA Polymerase, 2 mM (MgCl2), 4% DMSO, 0.2 mM dNTPs, 0.2 μM of each of the following primers: DiCSIP-Uni-Fwd, DiCSIP-Rev-AC, D-B, D-D and DiCSIP-Rev-F, and 2.5 ng gDNA template for *An. dirus*, *An. cracens*, and *An. baimaii*, or 0.1 ng pDNA template for *An. scanloni* and *An. nemophilous* ITS2 plasmids. Thermal cycling conditions consist of denaturation at 94°C for 2 min, followed by 35 cycles of denaturation at 94°C for 20 sec, annealing at 62 °C for 20 sec, and extension at 72°C for 1 min followed by final extension step at 72°C for 5 min.

The second PCR reaction is the SSP to differentiate *An. dirus* and *An. scanloni* by using a pair of DiCSIP*-*Fwd-C and DiCSIP-Rev-AC primers, which was conducted as described in the section 2.5.3.

### 2.7 Comparison of *An. dirus* complex identification by Dirus AS-PCR and DiCSIP in laboratory and field samples

The validation DiCSIP was performed using the conditions as described in the section 2.6. The assay was validated using Taq DNA polymerase from two different manufacturers (Promega Gotaq Flexi DNA polymerase or Invitrogen Taq DNA polymerase) in two laboratories, the National Center for Genetic Engineering and Biotechnology (BIOTEC) and Kasetsart University (KU). Thermal cycler used at BIOTEC was Biorad C1000 Touch Thermal Cycler, while the thermal cycler at KU was Bioer LifePro Thermal Cycler. Identification efficiency of DiCSIP was compared to the Dirus AS-PCR, which was conducted following a condition as described in the section 2.3.2 with a Ta of 55°C. Samples that resulted in amplicon sizes of approximately 520 bp in DiCSIP reaction were followed up by SSP to differentiate between *An. dirus* and *An. scanloni*.

## 3. Results

### 3.1 Dirus AS-PCR give inconsistent PCR amplification even after condition optimization

Genomic DNA of three known species within the *An. dirus* complex (*An. dirus*, *An. cracens*, *An. baimaii*) and plasmids harboring synthetic ITS2 constructs for *An. scanloni* and *An. nemophilous* were used as templates in the Dirus AS-PCR following the original protocol (9). The expected ITS2 amplicon sizes for the target species are 562 bp for *An. dirus*, 514 bp for *An. cracens*, 349 or 353 bp for *An. scanloni*, 306 bp for *An. baimaii*, and 223 bp for *An. nemophilous* (***Table 1***).

However, following the original Dirus AS-PCR, we encountered difficulties in species identification. Despite the presence of correct amplicons in *An. dirus* and *An. cracens* samples, several non-specific bands were present in these reactions. Moreover, we failed to obtain amplicons of the expected size for the other three species (***Figure 1A***). Additionally, the *An. scanloni* sample had an amplicon of the *An. dirus* expected size, which could lead to potential misidentification of *An. scanloni* as *An. dirus*. Given that the original Dirus AS-PCR protocol was developed over two decades ago, it is possible that variations in Taq polymerase manufacturers or thermal cyclers might be a cause for the PCR results differed from the original publication (9). Therefore, we conducted Dirus AS-PCR in different laboratories (BIOTEC or KU) and utilized Taq polymerases from different manufacturers (Invitrogen or Promega), yet the results showed non-specific and incorrect amplification (***Supplementary Figure S1A, B***).

**Figure 1.**
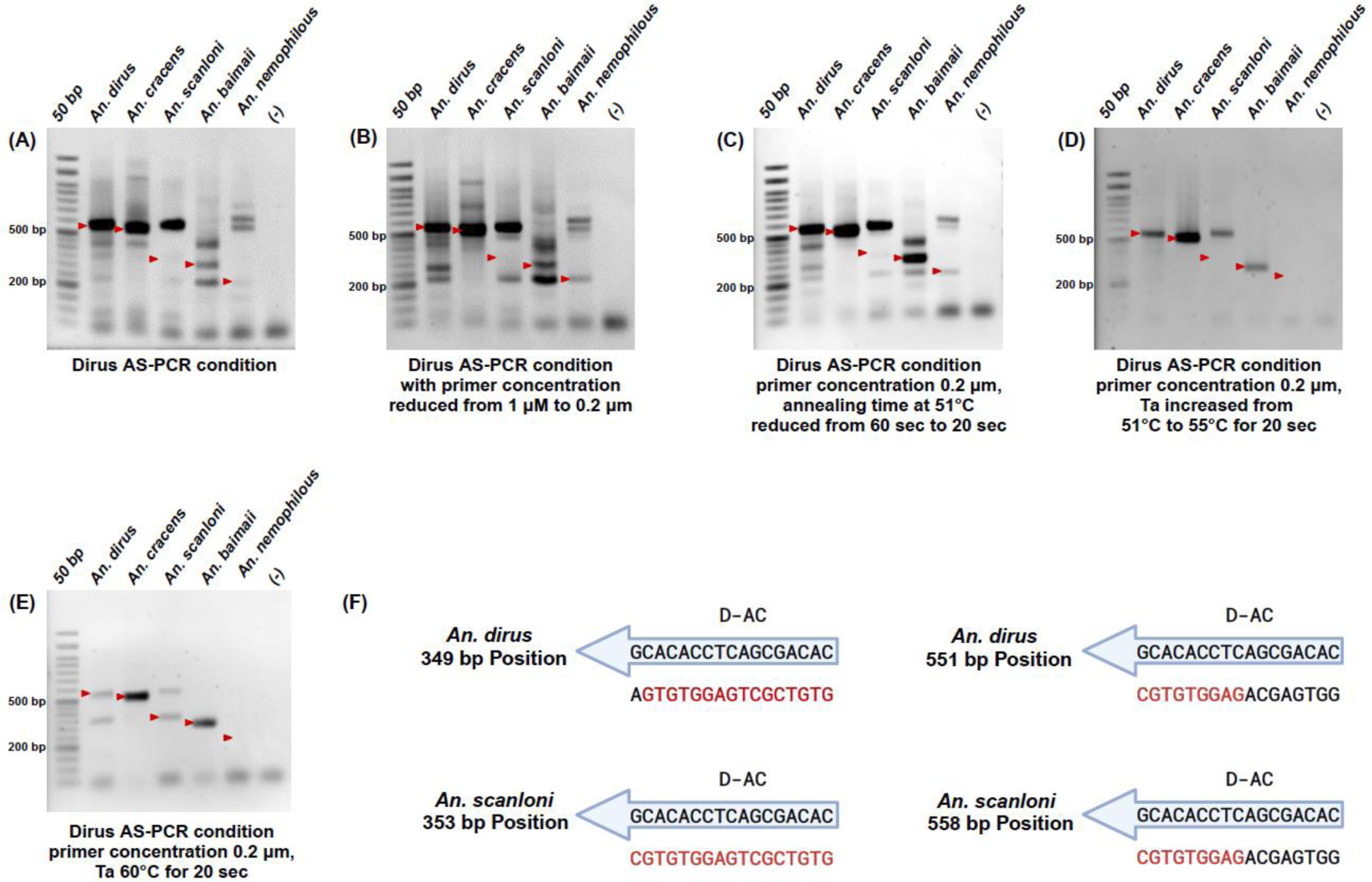
The original Dirus AS-PCR fails to correctly identify five sibling species of *An. dirus* complex present in the GMS even after optimization. Agarose gel electrophoresis of the amplicons from original PCR condition (A), a reduction of final primer concentration (B), reduction of incubation time (C), an increase of annealing temperature to 55°C (D), an increase of annealing temperature to 60°C (E). The red arrows indicate the expected sizes of PCR amplicons. (F) diagram demonstrating D-AC binding sites on *An. dirus* and *An. scanloni* ITS2 at 349-353 bp and 551-558 bp positions. Red letters indicate matching bases between D-AC and ITS2 templates.

To improve specificity of the assay, we modified the Dirus AS-PCR conditions. As an excessive primer concentration, especially in multiplex PCR, can lead to non-specific amplification, we first addressed this issue by reducing the final primer concentration from 1 µM to 0.2 µM, but the problems of non-specific and incorrect amplification still persisted (***Figure 1B***).

Then, we further optimized PCR condition by reducing incubation time as described in the method section 2.3.2. Although shorter PCR cycling time slightly improved the specificity of *An. dirus*, *An. cracens*, and *An. baimaii*, non-specific bands were still present under these conditions (***Figure 1C***). Furthermore, we attempted to reduce non-specific amplification by increasing the Ta from 51°C to 55°C resulting in correct amplifications for *An. dirus*, *An. cracens*, and *An. baimaii* (***Figure 1D***). While the Ta at 60°C resulted in correct amplifications of *An. cracens* and *An. baimaii* (***Figure 1E***), we obtained two bands of expected amplicons for *An. dirus and An. scanloni at* 562 and 350 bp for samples of both species. These results demonstrated that the original Dirus AS-PCR primers fail to correctly differentiate between *An. dirus* and *An. scanloni*, which might lead to misidentification between these two species. In addition, although the condition in ***Figure 1D*** demonstrated clean and correct amplicons for most species, inconsistencies were still observed with different enzyme manufacturers and thermal cyclers (***Supplementary Figure S2A, B, C, D***). We also found that the original Dirus AS-PCR primers can identify *An. dirus* and *An. scanloni* samples as *An. nemophilous* (***Supplementary Figure S2A, B***).

The results from this section confirm that the reverse primers for *An. cracens* (D-B) and *An. baimaii* (D-D) can correctly differentiate these two species after optimization of the PCR conditions, while highlighting the challenges encountered with reverse primers for the other three sibling species.

### 3.2 The original Dirus AS-PCR primers have multiple potential binding sites/amplicons

To identify the cause of difficulties in *An. dirus* molecular identification in the original Dirus AS-PCR, we conducted comprehensive bioinformatic analyses to identify potential off-target binding sites and non-specific amplification for each primer using primer tools in Benchling. Additionally, Primer-BLAST was used to evaluate off-target amplification of each primer pair including: i) potential off-target amplification within species, as well as ii) off-target cross-species amplification of ITS2 gene by each pair of universal forward and species-specific reverse primer pairs against nucleotide database of *Anopheles* genus (tax id:7164) and the results are summarized in the ***Table 1***.

#### 3.2.1 The D-U Dirus AS-PCR universal forward primer might cause difficulties in PCR amplification

Bioinformatic analysis of the D-U universal forward primer revealed its exceptionally high GC content at 88% and Tm at 68°C (***Table 1***) with multiple potential binding sites when the PCR conditions are less stringent (***Supplementary Figure S3***).

#### 3.2.2 *In silico* analyses reveal challenges in *An. dirus* and *An. scanloni* identification with the original D-U and D-AC primers

The *in silico* analyses using Benchling revealed that the D-AC primer has exact match to *An. scanloni* ITS2 that yield a single 349 or 353 bp amplicon when used with D-U, while *An. dirus* ITS2 has 3’ single nucleotide mismatch at the exact same location (***Figure 1F***, ***Table 1***, ***Supplementary Figure S4***). In addition to this location, the D-AC also has 9 bp out of 17 bp complementary leading to a 562 bp amplicon in both *An. dirus* and *An. scanloni* ITS2. Such design of D-AC allows the reverse primer to differentiate between *An. dirus* and *An. scanloni* because the presence of 3’ single nucleotide mismatch in *An. dirus* lowers the amplification efficiency (28), increasing the likelihood of 562 bp amplification in *An. dirus,* while *An. scanloni* yields 349 or 353 bp amplicon. The *in silico* analysis of D-AC explains the two bands observed in the PCR reactions since both amplicons are possible for *An. dirus* and *An. scanloni*.

Additionally, Primer-BLAST of the universal forward (D-U) and reverse primer (D-AC) did not provide any result for *An. scanloni* due to the unavailability of its ITS2 sequence in the NCBI database. Instead, the search returned several matches of 353 or 345 bp for *An. dirus* ITS2 (***Supplementary Figure S4A***), but not 562 bp due to insufficient partial match (only 9 out of 17 bp) on the latter location. Interestingly, while most *An. dirus* ITS2 sequences obtained by Primer-BLAST had this 3’ single nucleotide mismatch as expected, one of the *An. dirus* hit (OQ091691) had a complete match. This suggests a potential misidentification of *An. scanloni* as *An. dirus* in the NCBI database. The follow-up analysis by MSA between OQ091691, *An. dirus* and *An. scanloni* ITS2 from Walton et al. (1999) (9), and another *An. dirus* ITS2 from NCBI (MW647457) confirms that the OQ091691 should indeed be *An. scanloni* since the sequence lacks the deletion of CA repeats observed in *An. dirus* ITS2 (***Supplementary Figure S4B***).

#### 3.2.3 *In silico* analyses confirms high specificity of Dirus AS-PCR primers for *An. cracens* (D-U and D-B pair) and *An. baimaii* (D-U and D-D pair) identification

*In silico* binding site identification revealed that the *An. cracens* specific D-B primer has potential off-target binding sites albeit with much lower Tm (45.9°C) than the on-target binding site (56.2°C). The *An. baimaii* specific D-D primer had no potential off-target amplification (***Supplementary Figure S5***). Primer-BLAST of these two species only returned on-target ITS2 gene (***Table 1***).

#### 3.2.4 The off-target analysis of the original Dirus AS-PCR primers for *An. nemophilous* identification (D-U and D-F pair)

This set of primers also does not have cross or on-species off-target binding sites (***Supplementary Figure S5***). The Primer-BLAST did not return any result from *An. nemophilous* because the ITS2 sequence of this species is not available in the NCBI database (***Table 1***).

### 3.3. Development of new primers for Dirus Complex Species Identification PCR (DiCSIP)

The combination of experimental and *in silico* analyses above provide strong evidence that universal forward (D-U) and reverse primers (D-AC and D-F) were not efficient to identify *An. dirus*, *An. scanloni*, and *An. nemophilous*. Therefore, we designed a new set of primers for the identification of these three species, then used them in a multiplex PCR assay, hereinafter referred to as the Dirus Complex Species Identification PCR (DiCSIP)

#### 3.3.1 Design and validation of a new universal forward primer (DiCSIP-Uni-Fwd)

To design the DiCSIP-Uni-Fwd primer, MSA was conducted using ITS2 sequences from five species within the *An. dirus* complex (***Figure 2***, ***Supplementary Figure S6***), and the DiCSIP-Uni-Fwd primer was designed within the conserved region of the species. The properties of the new forward primer were greatly improved (57.14 %GC and 56.3°C Tm) compared to the original D-U primer (88.24 %GC and 68.22°C Tm) (***Table 2***). *In silico* potential binding site identification revealed only a single binding site on ITS2 in all five *An. dirus* complex species (***Supplementary Figure S7)*** suggesting improved specificity compared to the previous primer (D-U).

**Figure 2.**
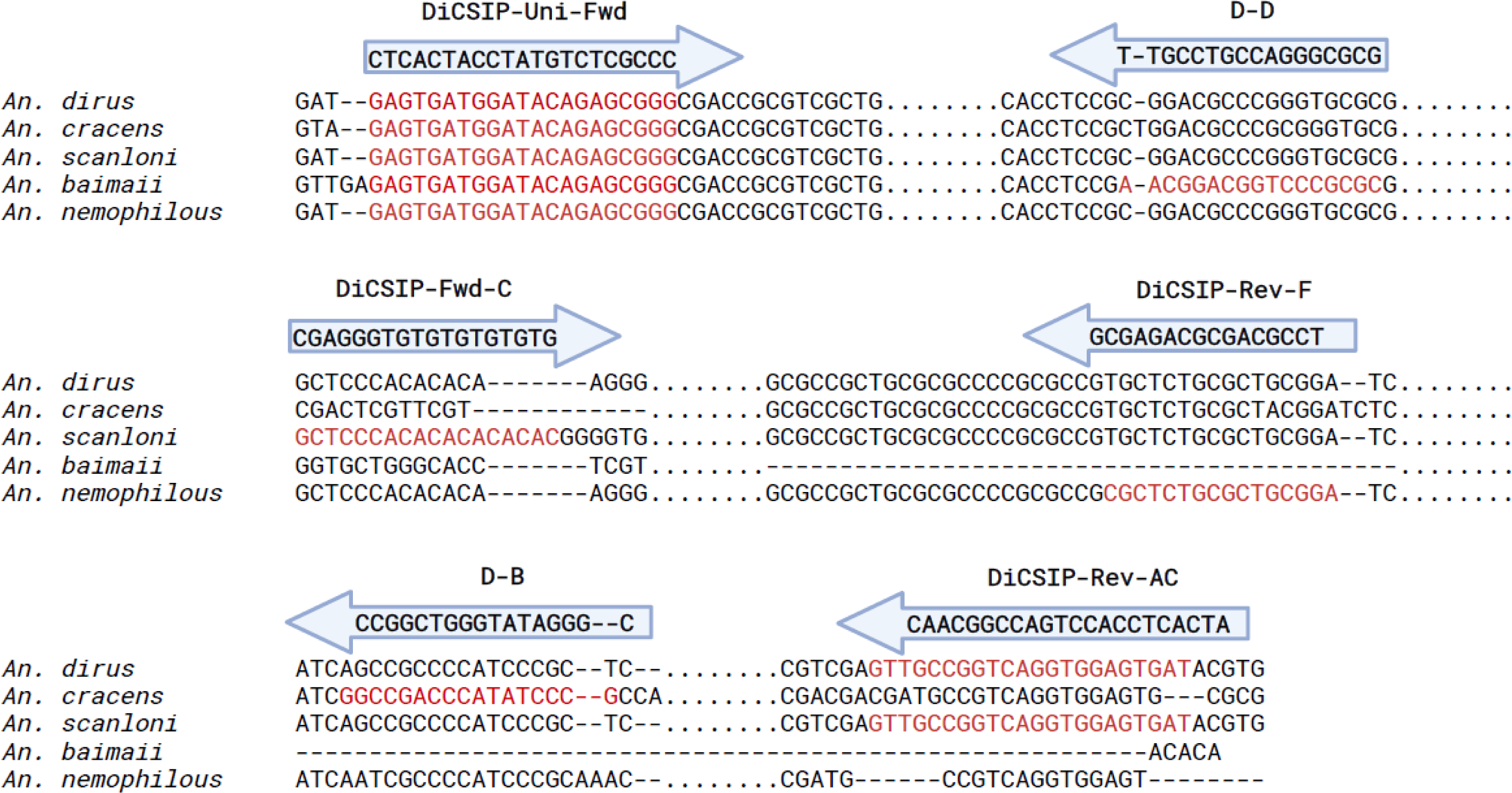
Multiple sequence alignment of ITS2 sequences of five *An. dirus* complex sibling species and binding sites of DiCSIP primers. The sequences of *irus* (Accession: MW647457.1), *An. cracens* (Accession: MG008574.1) and *An. baimaii* (Accession: MN152993.1) were retrieved from NCBI, while those of *An. oni* and *An. nemophilous* were obtained from a publish article (9). The binding sites of each primer were highlighted in red. Dots (.) indicate shortened ences, while dashes (-) indicate base insertion/deletion. The full alignment can be found in the ***Supplementary Figure S6***.

Following the *in silico* design of DiCSIP-Uni-Fwd, we conducted a PCR to assess whether this primer could enhance the specificity of the Dirus AS-PCR. In this experiment, the original universal forward primer D-U was replaced by the new forward primer DiCSIP-Uni-Fwd, while keeping all the original reverse primers. The PCR was conducted with a final primer concentration of 0.2 μM each (DiCSIP-Uni-Fwd, D-AC, D-B, D-D, and D-F), and the same thermal cycling condition as ***Figure 1E***. The DiCSIP-Uni-Fwd was 79 bp downstream from the original forward primer. Thus, the expected amplicons were 483-489 bp for *An. dirus,* 435 bp for *An. cracens*, 287-294 bp for *An. scanloni*, 227 bp for *An. baimaii*, and 144 bp for *An. nemophilous*. We found that the replacement of only the forward primer could not solve the problem with *An. dirus*, *An. scanloni,* and *An. nemophilous* samples (***Figure 3***). Therefore, a new set of reverse primers were needed to improve PCR species identification.

**Figure 3.**
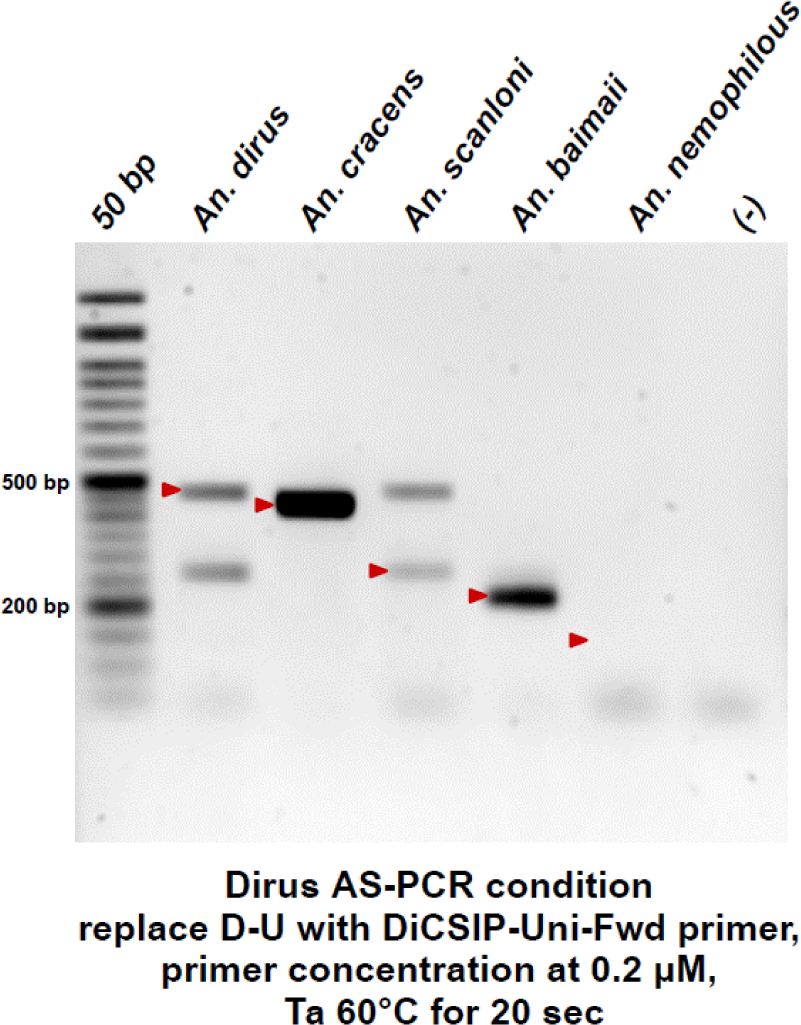
The replacement of the universal forward primer alone does not solve the problem of Dirus AS-PCR. The PCR was conducted using the DiCSIP-Uni-Fwd primer and all original reverse primers (DiCSIP-Uni-Fwd, D-AC, D-B, D-D, and D-F) at final primer concentration of 0.2 μM each. The red arrow mark indicates the expected sizes of PCR amplicons.

#### 3.3.2. Design and validation of new DiCSIP primers for *An. dirus* and *An. scanloni* identification

Due to high sequence similarity between *An. dirus* and *An. scanloni* ITS2, designing a new reverse primer that targets only *An. dirus* or *An. scanloni* ITS2 at different binding sites was impossible. Therefore, we designed two sets of reverse primers, one targeting both species (DiCSIP-Rev-AC) and another one targeting only *An. scanloni* (DiCSIP-Rev-C2 to C6). With these two primers in a multiplex reaction, it was expected that *An. dirus* will have one amplicon from DiCSIP-Rev-AC primer, while *An. scanloni* will have additional amplicon from DiCSIP-Rev-AC and DiCSIP-Rev-C primers.

The DiCSIP-Rev-AC primer has a GC content of 59.09% and a Tm of 60.88°C. The amplicon size from DiCSIP-Uni-Fwd and DiCSIP-Rev-AC primers was expected to be 521 bp for *An. dirus* and 528 bp for *An. scanloni* (***Table 2***). The analysis revealed high specificity with no off-target binding sites (***Supplementary Figure S8***). The validation by single-plex PCR containing the DiCSIP-Uni-Fwd and DiCSIP-Rev-AC primers resulted in specific amplicons of approximately 520 bp for both *An. dirus* and *An. scanloni* (***Figure* 4*A***, ***B***).

**Figure 4.**
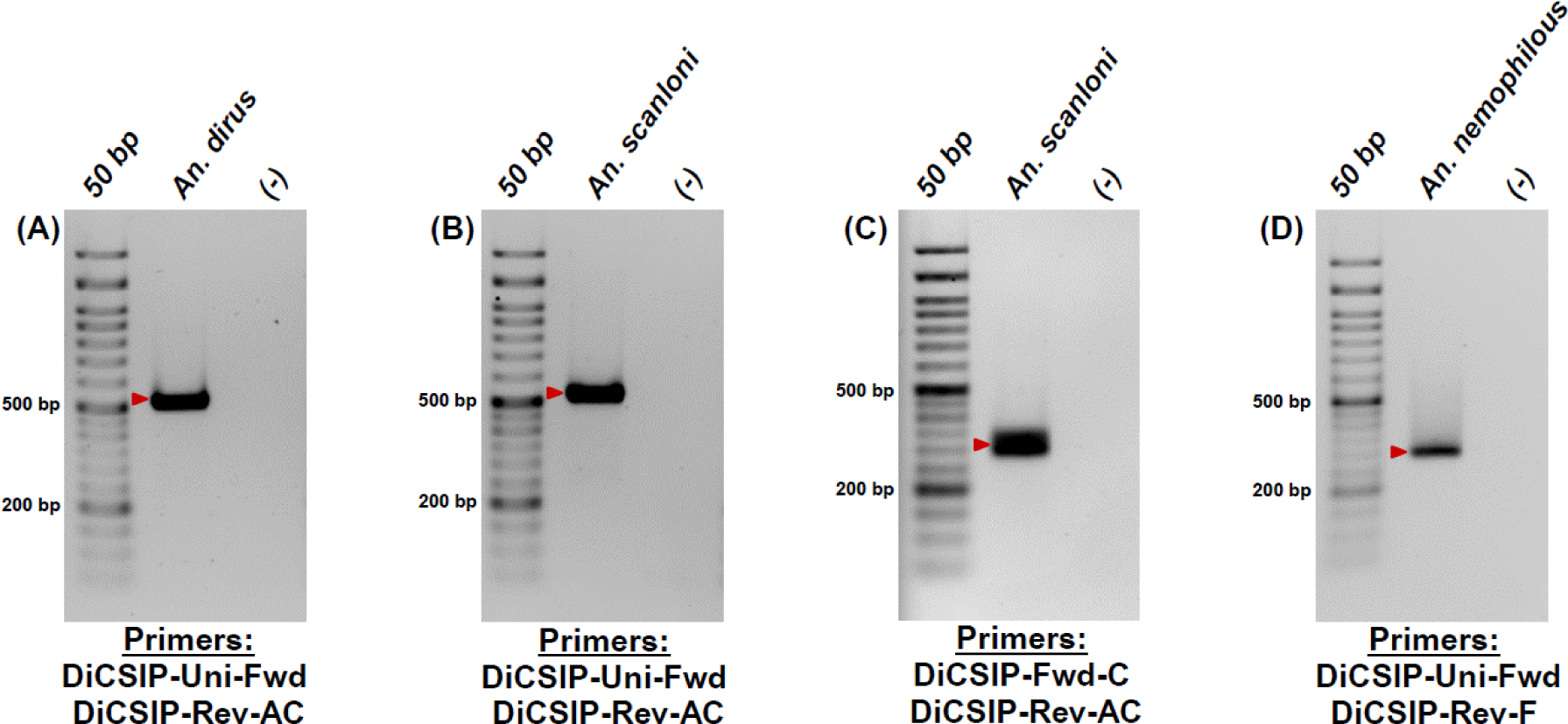
Single-plex PCR demonstrates high specificity of the new primers for *An. dirus, An. scanloni,* and *An. nemophilous* identifications. Validation of the new *An. dirus/scanloni* specific reverse primers to identify *An. dirus* (A) and *An. scanloni* (B). Validation of the new *An. scanloni* specific forward primer to identify *An. scanloni* (C). Validation of the new *An. nemophilous* specific reverse primer to identify *An. nemophilous* (D). The red arrows indicate the expected sizes of PCR amplicon for each species.

Six reverse primers were designed to target the *An. scanloni* ITS2 and differentiate this species from *An. dirus* with primer properties in the range of 57.14 – 68.75% GC and 55.3 – 61.0°C Tm (***Supplementary Table S1***). *In silico* analyses of amplicons between DiCSIP-Uni-Fwd and the six new *An. scanloni* reverse primers predicted amplicon sizes ranging from 246 to 272 bp (***Supplementary Table S1, Supplementary Figure S9***). However, the PCR validation demonstrated that none of the new *An. scanloni* reverse primers yielded correct amplicons (***Supplementary Figure S9***).

As no alternative region was available for designing *An. scanloni* ITS2 specific reverse primers, we then changed the strategy for *An. scanloni*-specific forward primer (DiCSIP-Fwd-C) (***Supplementary Figure S10***). Validation by PCR with the DiCSIP-Fwd-C and DiCSIP-Rev-AC primers using the *An. scanloni* ITS2 plasmid as a template showed a clean amplification at 300 bp (***Figure 4C***).

#### 3.3.3. Design and validation of new DiCSIP primer for *An. nemophilous* identification

A new reverse primer specifically targeting *An. nemophilous* ITS2 (DiCSIP-Rev-F) was designed with primer properties of 75% GC and 60.27°C Tm. This primer has an exact match binding site to *An. nemophilous* ITS2 (***Supplementary Figure S11***), resulting in an expected amplicon size of 223 bp (***Table 2***). Although the *in silico* analysis suggested potential off-target binding sites of DiCSIP-Rev-F primer on *An. nemophilous* ITS2, this is the only available region to design *An. nemophilous* specific primer (***Supplementary Figure S11***). Nevertheless, single-plex PCR validation with DiCSIP-Uni-Fwd and DiCSIP-Rev-F primers with *An. nemophilous* plasmid exhibited a single 223 bp amplicon (***Figure 4D***).

### 3.4 The new multiplex DiCSIP improves identification of *An. dirus* complex sibling species

After successfully validating the specificity of the new primers in single-plex PCR, we combined all six primers including two forward primers (DiCSIP-Uni-Fwd and DiCSIP-Fwd-C) and four reverse primers (DiCSIP-Rev-AC, D-B, D-D, and DiCSIP-Rev-F) in a multiplex PCR at final concentration of 0.2 μM each. The amplification was conducted as described in section 2.5.1. The amplicon sizes for each species were listed in ***Table 2***. Unexpectedly, the multiplex PCR with two forward and four reverse primers in one reaction (***Figure 5A***) could correctly identify only *An. dirus* (521 bp), *An. cracens* (435 bp). While two amplicons were expected from *An. scanloni* sample at 521 and 300 bp, the actual PCR amplification only resulted in a single band at 521 bp suggesting that the multiplex reaction might interfere with the binding of *An. scanloni*-specific DiCSIP-Fwd-C primer. In addition to the on-target amplification for *An. baimaii* (225 bp) and *An. nemophilous* (305 bp), these samples also exhibited a fainter off-target band at approximately 560 bp and 520 bp for these two species, respectively. These results suggested that DiCSIP-Fwd-C primer might interfere with other primers making it challenging to accurately differentiate sibling species within the *An. dirus* complex using one multiplex PCR reaction. Therefore, separate PCR reactions are needed to ensure precise species identification, the first reaction with the DiCSIP-Uni-Fwd forward primer and four reverse primers (DiCSIP-Rev-AC, D-B, D-D, and DiCSIP-Rev-F) to identify *An. dirus*/*scanloni*, *An. cracens*, *An. baimaii*, and *An. nemophilous*, based on their respective amplicon sizes. The second Scanloni-specific PCR (SSP) assay aims to differentiate *An. dirus* and *An. scanloni* by using a pair of DiCSIP*-*Fwd-C and DiCSIP-Rev-AC primers with expected size at 300 bp.

**Figure 5.**
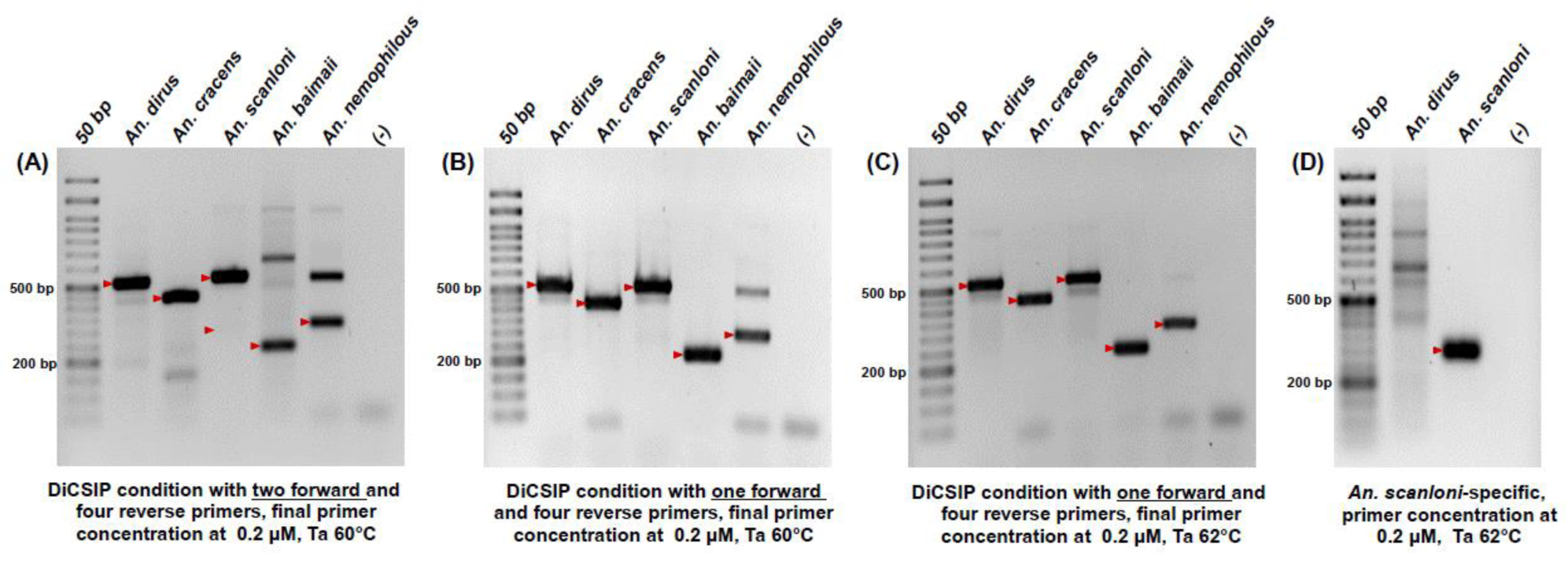
DiCSIP correctly identifies five sibling species of the *An. dirus* complex. (A) DiCSIP reaction containing two forward (DiCSIP-Uni-Fwd and DiCSIP-Fwd-C) and four reverse primers (DiCSIP-Rev-AC, D-B, D-D and DiCSIP-Rev-F). (B) DiCSIP reaction containing one forward and four reverse primers DiCSIP-Uni-Fwd, DiCSIP-Rev-AC, D-B, D-D and DiCSIP-Rev-F. (C) DiCSIP reaction increasing annealing temperature from 60°C to 62°C using one forward and four reverse primers DiCSIP-Uni-Fwd, DiCSIP-Rev-AC, D-B, D-D and DiCSIP-Rev-F. (D) SSP reaction containing DiCSIP-Fwd-C and DiCSIP-Rev-AC primers correctly differentiate *An. scanloni* from *An. dirus*. The red arrow mark indicates the expected sizes of PCR amplicons.

The DiCSIP reaction was initially validated using a Ta of 60°C, which accurately identify all five species, albeit with a fainter off-target band at 520 bp for *An. nemophilous* (***Figure 5B***) The non-specific band in *An. nemophilous* sample could be eliminated by increasing the Ta to 62°C (***Figure 5C***). The differentiation between *An. dirus* and *An. scanloni* samples by SSP was validated with a Ta of 62°C, resulting in a correct amplicon at 300 bp for the *An. scanloni* sample (***Figure 5D***). Although off-target amplification appeared as a smear larger than 300 bp for the *An. dirus* sample, the specificity could not be further improved as this is the only region available for designing an *An. scanloni*-specific primer.

### 3.4 The DiCSIP improves sensitivity of *An. dirus* species complex identification

Primers with a wide operational range and high sensitivity are crucial for reproducibility, especially across different laboratories. Since *An. dirus* was the most problematic sample in the original PCR, we compared operational range of the Dirus AS-PCR and DiCSIP by using varying concentrations of *An. dirus* gDNA template, ranging from 10 ng to 0.0001 ng/12.5 µL reaction. The results showed that the DiCSIP consistently and accurately produced correct amplicon across a wide range of template concentrations from 10 ng to 0.01 ng (***Figure 6A***). In contrast, the original Dirus AS-PCR had a much more limited working range for DNA templates, spanning from 10 ng to 1 ng (***Figure 6A***). Additionally, the intensity of the amplicon from reaction with 10 ng template using Dirus AS-PCR was fainter than those from 0.1 ng template using DiCSIP. These results demonstrated at least a 100-fold improvement in sensitivity by DiCSIP.

**Figure 6.**
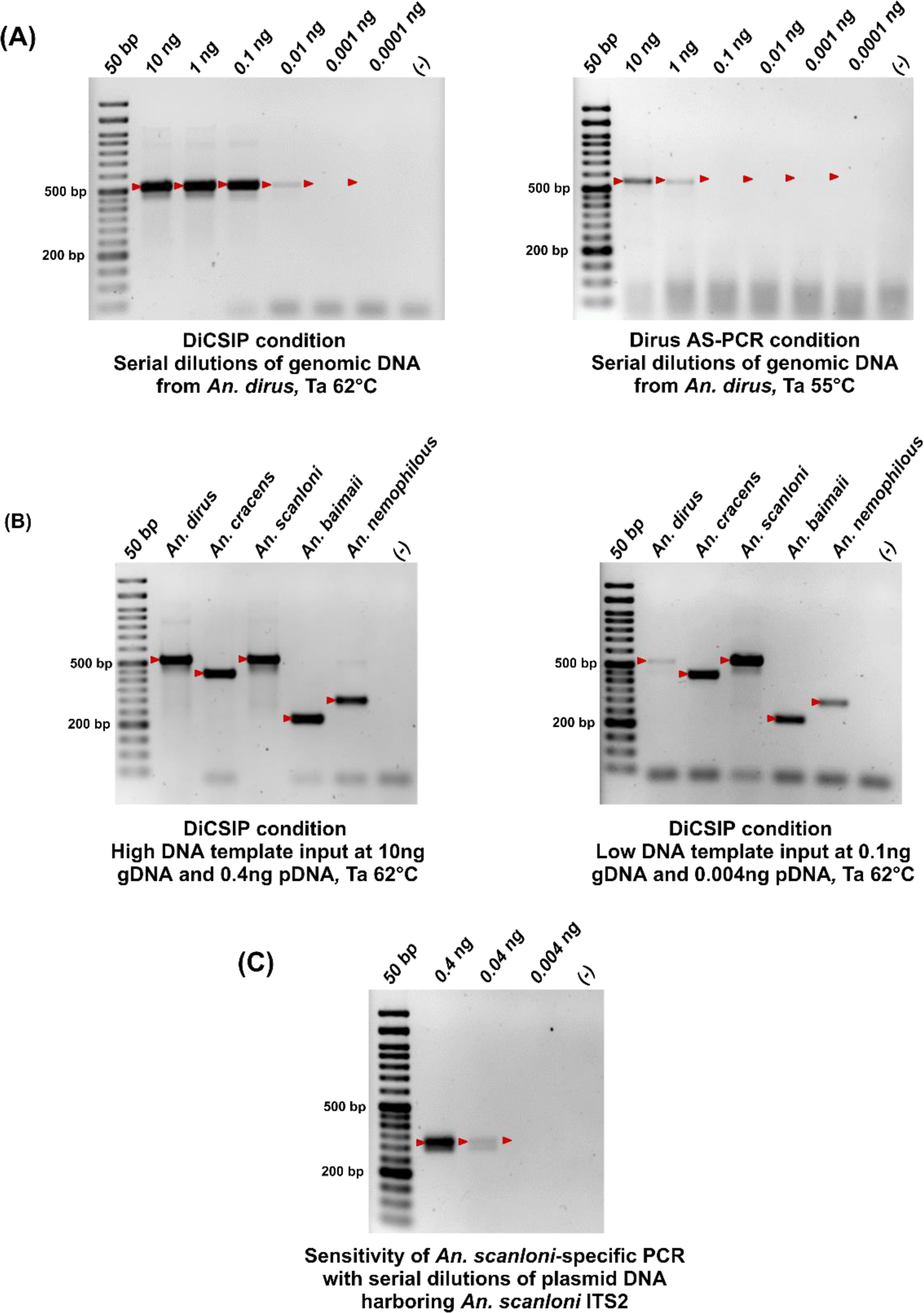
DiCSIP has higher sensitivity and wider operational range than Dirus AS-PCR. (A) Sensitivity of DiCSIP and Dirus AS-PCR was determined using *An. dirus* gDNA ranging from 0.0001-10 ng per reaction. (B) Identification of five *An. dirus* complex sibling species with high (10 ng gDNA or 0.4 ng pDNA) and low quantity (0.1 ng gDNA or 0.004 ng pDNA) of template by DiCSIP. (C) Operational range of SSP for *An. scanloni* identification with 0.004-0.4 ng of *An. scanloni* ITS2 plasmid DNA. The red arrows indicate the expected sizes of PCR amplicons.

To expand the observation to other sibling species of the *An. dirus* complex, gDNA templates at 10 ng and 0.1 ng or pDNA templates at 0.4 and 0.004 ng were used in DiCSIP to evaluate the operational range for species identification. The results in ***Figure 6B*** confirmed that DiCSIP effectively identify five sibling species of the *An. dirus* complex both at high and low template quantities per 12.5 µL reaction.

Similarly, operational range of the SSP was assessed with pDNA template, ranging from 0.4 ng to 0.004 ng/12.5 µL reaction. The results demonstrated strong amplification at 0.4 ng, while the amplicon at 0.04 ng had low intensity (***Figure 6C***).

### 3.5 DiCSIP outperforms Dirus AS-PCR in *An. dirus* complex identification

To demonstrate the efficiency of DiCSIP, we compared species identification between DiCSIP and the original Dirus AS-PCR on laboratory and field specimens that were morphologically identified as *An. dirus*. These tests were carried out at two different laboratories, BIOTEC and KU.

The DiCSIP consistently and correctly identified all laboratory specimens of *An. dirus* and *An. cracens* in both laboratories. However, the original Dirus AS-PCR yielded contradictory results between laboratories for *An. dirus*, specifically, the original Dirus AS-PCR correctly identified *An. dirus* specimens at KU, but the same samples were incorrectly identified as *An. nemophilous* at BIOTEC (***Table 3***). In contrast, all *An. cracens* laboratory specimens were correctly identified by assays in both laboratories (***Table 3***).

**Table 3.** Summary of *An. dirus* complex species identification by DiCSIP and Dirus AS-PCR in laboratory and field-collected specimens.

| Laboratory | Insectary/Locality | *An. dirus* |  | *An. cracens* |  | *An. scanloni* |  | *An. baimaii* |  | *An. nemophilous* |  |
| --- | --- | --- | --- | --- | --- | --- | --- | --- | --- | --- | --- |
|  |  | Original<br>Dirus AS-PCR | DiCSIP | Original<br>Dirus AS-PCR | DiCSIP | Original<br>Dirus AS-PCR | DiCSIP | Original<br>Dirus AS-PCR | DiCSIP | Original<br>Dirus AS-PCR | DiCSIP |
| KU | *An. dirus* laboratory colony | 10 | 10 | - | - | - | - | - | - | - | - |
|  | *An. cracens* laboratory colony | - | - | 10 | 10 | - | - | - | - | - | - |
|  | Kanchanaburi Province | 4 | 4 | - | - | - | - | 6 | 6 | - | - |
|  | Prachinburi Province | 9 | 9 | - | - | - | - | 1 | 1 | - | - |
|  | Ranong Province | - | - | - | - | - | - | 10 | 10 | - | - |
|  | Sisaket Province | 10 | 10 | - | - | - | - | - | - | - | - |
| BIOTEC | *An. dirus* laboratory colony | - | 10 | - | - | - | - | - | - | 10* | - |
|  | *An. cracens* laboratory colony | - | - | 10 | 10 | - | - | - | - | - | - |
|  | Kanchanaburi Province | - | 4 | - | - | - | - | 6 | 6 | 4* | - |
|  | Prachinburi Province | - | 9 | - | - | - | - | 1 | 1 | 9* | - |
|  | Ranong Province | - | - | - | - | - | - | 10 | 10 | - | - |
|  | Sisaket Province | - | 10 | - | - | - | - | - | - | 10* | - |

\*The samples were identified as *An. dirus* by the DiCSIP

Next, the validation of the primers was extended to field-collected specimens morphologically identified as *An. dirus* complex, with ten specimens from each of four collection sites in Thailand: Kanchanaburi, Prachinburi, Ranong, and Sisaket Provinces. We found that both the DiCSIP and the original Dirus AS-PCR provided the same identification results for all *An. cracens* and *An. baimaii* samples regardless of collection sites and laboratory conducting the identification (***Table 3***). However, while the DiCSIP correctly identify field-derived *An. dirus* regardless of laboratory conducted the assay, the original Dirus AS-PCR correctly identified *An. dirus* specimens at KU but the same samples were incorrectly identified as *An. nemophilous* at BIOTEC (***Table 3***).

## 4. Discussion

Members of *An. dirus* complex present in the GMS, including Thailand, have been incriminated as vectors for human malaria, with two species, *An. dirus* and *An. baimaii*, recognized as main malaria vectors (22–25, 29) while the three other species are recognized as either secondary/incidental vectors (*An. cracens*, *An. scanloni*), or even non-vector (*An. nemophilous*) (20). Six of the eight sibling species of the *An. dirus* complex share overlapping spatial distributions (sympatry) but exhibit distinct behaviors and vector competence. Given these complexities, precise identification of these sibling species is of great importance to fully comprehend and accurately define local malaria transmission dynamics. This is critical for the implementation of appropriate vector control strategies maximizing cost-effectiveness and resource utilization, especially as Thailand aims to achieve malaria elimination by 2024 and the GMS aims for elimination by 2030 (30). Even after elimination, accurate vector surveillance is important to monitor potential re-introduction of malaria to the GMS. The accurate surveillance is also important in our combat with the emerging problem of zoonotic transmission of simian malaria.

Primer-BLAST results in the current study raised our concern for misidentification of sibling species of the *An. dirus* complex. We found that some of the *An. dirus* ITS2 in the NCBI was misidentified, and should be *An. scanloni*. Without *An. scanloni* ITS2 available in the database, the future BLAST search of *An. scanloni* ITS2 sequence will return that *An. dirus* accession and might lead to false identification (***Supplementary Figure S4*)**. Without careful analysis, subsequent study of the misidentified samples might lead to false advancements in research on *An. dirus* complex, as well as incorrect information on the geographical distribution of each species. Not only misidentification between *An. dirus* and *An. scanloni*, but we also found misidentification of *An. dirus* as *An. baimaii* in several NCBI entries (***Supplementary Figure S12B***). Therefore, a comprehensive revision of *An. dirus* complex ITS2 in NCBI is needed to improve accuracy of future species identification.

The original molecular identification assay, Dirus AS-PCR, was developed over two decades ago to differentiate five sibling species of the *An. dirus* complex found in the GMS (9). Since the development of the Dirus AS-PCR for *An. dirus* complex, several articles demonstrated adaptations of this protocol to identify the sibling species (26, 31–40). Despite the need of accurate entomological information, we found that the existing Dirus AS-PCR (9) yielded inconsistent results in our laboratory compared to the original article. In addition to results demonstrated in our study, inaccurate identification was also described by Monthatong (26), whom demonstrated challenges in correct identification of *An. dirus* and *An. scanloni*. Since the ITS2 region has secondary structures and complementary sequences throughout the gene, the development of PCR diagnostic based on ITS2 can be difficult, and slight changes in reaction components, conditions and equipment may affect the accuracy of identification. Indeed, our study demonstrated that the difference in thermal cycler used for PCR can cause misidentification.

In addition to the differences in technical aspects, we should also consider a biological aspect that the sequences used for primer design in the previous study might not adequately represent the full range of genetic diversity within these species, or the ITS2 sequences of the *An. dirus* complex may have evolved over time. Indeed, we found that the *An. dirus* ITS2 sequence described in Walton et al., 1999 was slightly different from the more recent *An. dirus* ITS2 (***Supplementary Figure S4B***). Although these polymorphisms might not directly be located on primer binding sites, they might cause changes in thermodynamics of template DNA, thus resulting in off-target or inefficient amplification. Another evidence of nucleotide polymorphisms that affect Dirus AS-PCR primer binding is demonstrated in the ***Supplementary Figure S12A***. Primer-BLAST results revealed that the binding site of D-AC primer on *An. dirus* ITS2 sequences misidentified as *An. baimaii* contains additional mismatch in addition to the 3’ single nucleotide mismatch.

Another important factor influencing accuracy of the Dirus AS-PCR primers is the handling during reaction setup. Taq polymerase is active at room temperature and can cause unintended amplification if the reaction is not kept on ice during the preparation. We found that the reaction set up had to be conducted on ice all the time to avoid non-specific amplification. Hot-start enzymes can be a viable alternative to reduce unintended amplification, albeit with higher cost per reaction.

We found that the different primers of the previous set of Dirus AS-PCR had high GC content, which leads to high Tm. The recommended Ta of 51°C in the protocol was much lower than Tm of the original primers, thus might increase non-specific binding. After optimization of the PCR conditions by increasing the temperature at the annealing step, coupled with reducing primer concentrations, improvement in PCR results was obtained. Nevertheless, the specificity of the primers still hinders the use of this protocol to identify species within the *An. dirus* complex.

A thorough *in silico* analysis pinpointed problematic primers in the original Dirus AS-PCR and allowed the design of DiCSIP primers, which improved specificity, operational range, and sensitivity enabling the accurate identification of all five member species of the GMS. Although requiring two separate reactions to differentiate all five species, it is important to note that even with the original Dirus AS-PCR, a second SSP reaction is required to differentiate *An. dirus* and *An. scanloni*, as our investigation demonstrated that the Dirus AS-PCR had difficulty discriminating between these two species. Two-step PCR process has also been used in the development of the multiplex PCR for identifying five sibling species of the *An. barbirostris* complex (10).

Validation using laboratory and field samples demonstrated that DiCSIP offers correct identification even in different laboratories, using different Taq reagents and thermal cyclers, which show the reproducibility of the new primer set. The wide operational range and high specificity allow us to use the DNAzol direct reagent to process mosquito samples and use them directly in the PCR reaction without the need for DNA extraction, thereby saving both time and cost in sample processing.

While it would have been ideal to include, in the development of DiCSIP, *An. aff. takasagoensis,* the sixth member of *An. dirus* complex found in the GMS, specifically in a restricted area of northern Vietnam (21), unfortunately its ITS2 sequence is not available in the NCBI database. This extends to *An. scanloni* and *An. nemophilous,* which also lack ITS2 sequence information and only those in Walton et al., 1999 (9) are available. The lack of molecular information on *An. dirus* complex members underscores a significant knowledge gap in genetic information of these species. This lack of recent molecular and population genetic information can hinder efforts for effective and sustainable malaria vector control as it is impossible to evaluate if the tools for species identification are still efficient.

Taken together, this study addresses significant challenges in *An. dirus* complex identification by developing the DiCSIP assay, which improves the specificity and efficiency of species identification. The new primer set provides a valuable tool for accurate entomological surveys, supporting efficient vector control measures to reduce malaria transmission and prevent the re-introduction of the disease in the GMS region. The study emphasizes the importance of molecular information to develop tools for reliable identification of sibling species of *Anopheles* complexes and improve malaria control strategies.

## Author contributions

<u>Conceptualization:</u> Manop Saeung, Natapong Jupatanakul

<u>Formal analysis:</u> Manop Saeung, Natapong Jupatanakul

<u>Funding acquisition:</u> Sylvie Manguin, Theeraphap Chareonviriyaphap, Natapong Jupatanakul

<u>Investigation:</u> Manop Saeung, Jutharat Pengon, Chatpong Pethrak, Sarunya Thaiudomsap, Suthat Lhaosudto, Natapong Jupatanakul

<u>Methodology:</u> Manop Saeung, Jutharat Pengon, Natapong Jupatanakul

<u>Project administration</u>: Natapong Jupatanakul.

<u>Resources:</u> Atiporn Saeung, Sylvie Manguin, Theeraphap Chareonviriyaphap, Natapong Jupatanakul

<u>Software:</u> Manop Saeung, Natapong Jupatanakul

<u>Supervision:</u> Sylvie Manguin, Theeraphap Chareonviriyaphap, Natapong Jupatanakul

<u>Validation:</u> Manop Saeung, Jutharat Pengon, Natapong Jupatanakul

<u>Visualization:</u> Manop Saeung, Natapong Jupatanakul

<u>Writing –original draft:</u> Manop Saeung, Natapong Jupatanakul

<u>Writing –review & editing:</u> Manop Saeung, Jutharat Pengon, Chatpong Pethrak, Sarunya Thaiudomsap, Suthat Lhaosudto, Atiporn Saeung, Sylvie Manguin, Theeraphap Chareonviriyaphap, Natapong Jupatanakul

