## Supplementary Table S1 for "Dirus Complex Species Identification PCR (DiCSIP) improves identification of *Anopheles dirus* complex from Greater Mekong Subregion"

**Supplementary Table S1**. Expected amplicon sizes of six An. scanloni-specific reverse primers when using with DiCSIP-Uni-Fwd primer

| **Primer** | **Direction** | **Sequence (5′―> 3′)** | **Length** | **Tm (°C)** | **%CG** | **Specificity within Dirus complex** | |
| --- | --- | --- | --- | --- | --- | --- | --- |
|  |  |  |  |  |  | **Expected size  (Product size)** | **Primer-BLAST Results  (Product size)** |
| DiCSIP-Uni-Fwd | Forward | GAGTGATGGATACAGAGCGGG | 21 bp | 56.3 | 57.14 | - | - |
| DiCSIP-Rev-C2V1 | Reverse | CGCACACACCCCGTGTGT | 18 bp | 60.0 | 66.67 | *An. scanloni* (259 bp) | No ITS2 sequence in the database |
| DiCSIP-Rev-C2V2 | Reverse | CGCACACACCCCTGTGTGT | 19 bp | 59.6 | 63.16 | *An. scanloni* (258 bp) | No ITS2 sequence in the database |
| DiCSIP-Rev-C3 | Reverse | GCACACACCCCGTGTGTG | 18 bp | 58.4 | 66.67 | *An. scanloni* (257 bp) | No ITS2 sequence in the database |
| DiCSIP-Rev-C4 | Reverse | CACACGCACACACCCC | 16 bp | 55.3 | 68.75 | *An. scanloni* (246 bp) | No ITS2 sequence in the database |
| DiCSIP-Rev-C5 | Reverse | CGCACACACCCCGTGTGTG | 19 bp | 61.0 | 68.42 | *An. scanloni* (258 bp) | No ITS2 sequence in the database |
| DiCSIP-Rev-C6 | Reverse | ACAGCGACTCCACACGC | 17 bp | 57.0 | 64.71 | *An. scanloni* (272 bp) | No ITS2 sequence in the database |
