## Supplementary Figure S1-S12 for "Dirus Complex Species Identification PCR (DiCSIP) improves identification of *Anopheles dirus* complex from Greater Mekong Subregion"

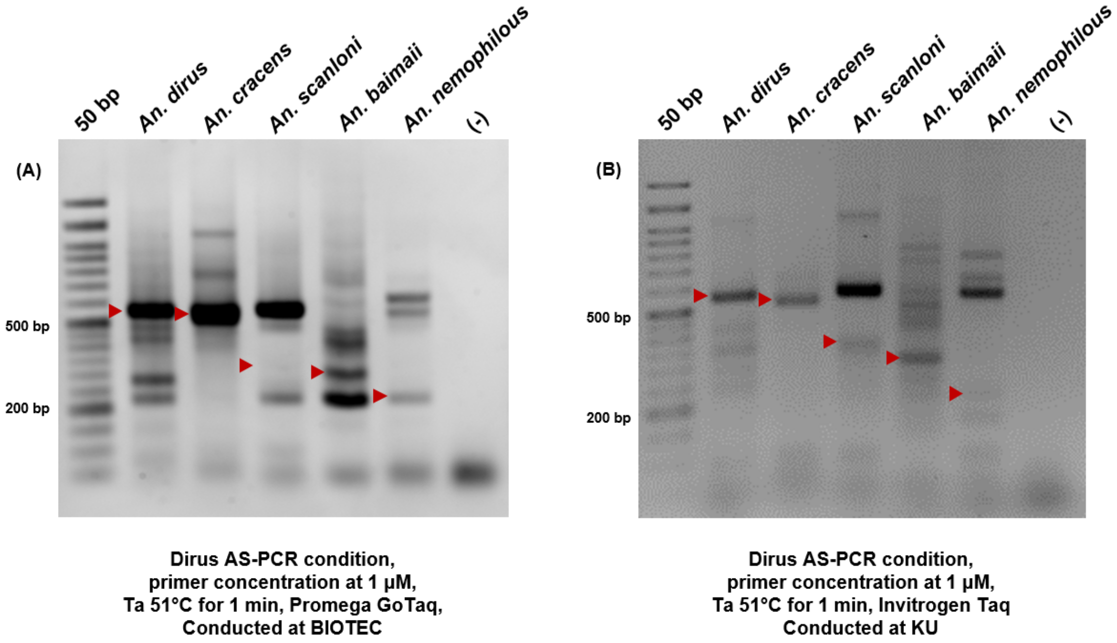


***Supplementary Figure S1*. The original Dirus AS-PCR assay fails to identify members of *An. dirus* complex in different laboratory.** Dirus AS-PCR was conducted following the exact conditions as described by Walton et al. (1999) in two laboratories to identify *An. dirus*, *An. cracens, An. scanloni*, *An. baimaii*, and *An. nemophilous*. (A) AS-PCR conducted at BIOTEC using Promega GoTaq Flexi DNA Polymerase with BioRad C1000 Touch Thermal Cycler. (B) AS-PCR performed at KU using Invitrogen Taq DNA polymerase with Bioer LifePro Thermal Cycler. The red arrow mark indicates the expected sizes of PCR amplicons.

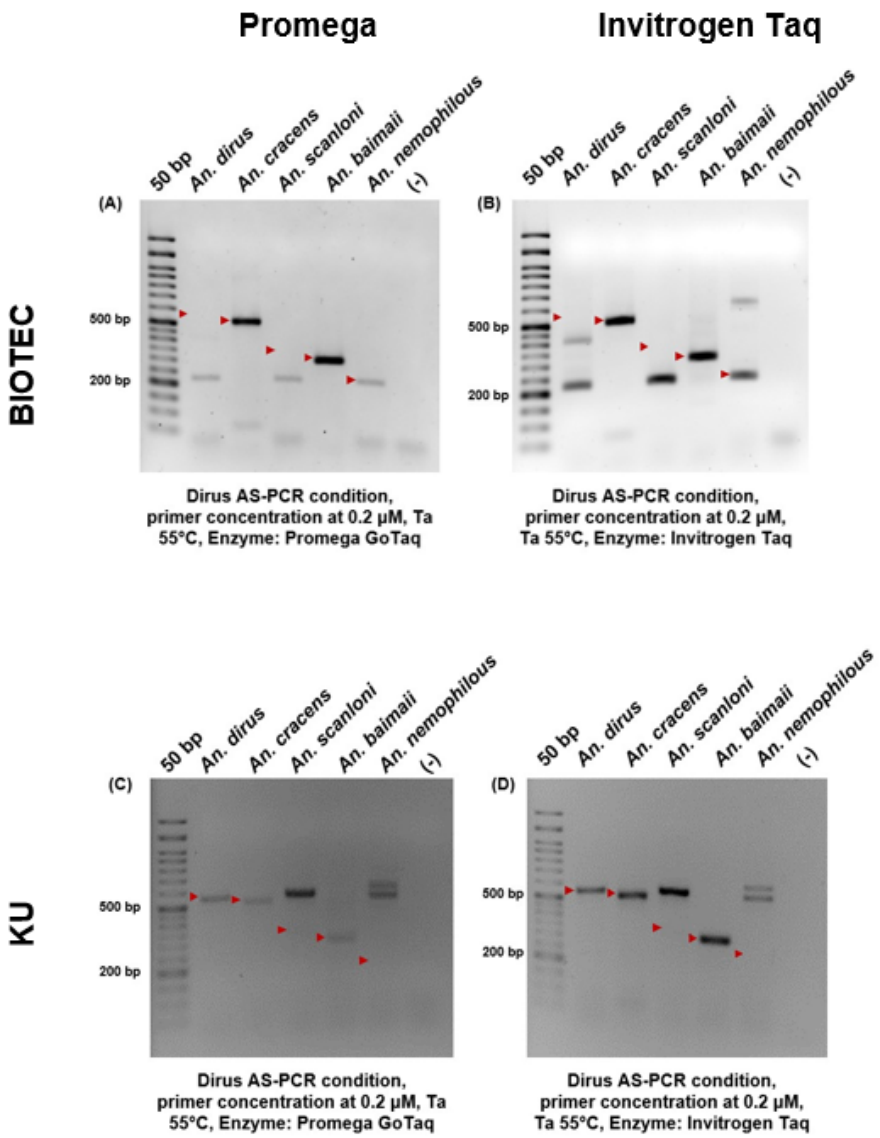


***Supplementary Figure S2.*** **The Dirus AS-PCR assay fails to consistently identify members of *An. dirus* complex even after condition optimization.** The Dirus AS- PCR was conducted using an optimized condition with different laboratories, Taq enzyme from different manufacturers and different thermal cyclers. The Dirus AS-PCR was performed at BIOTEC using Promega GoTaq Flexi DNA polymerase (A) and Invitrogen Taq DNA polymerase (B) with C1000 Touch Thermal Cycler, and at KU using Promega GoTaq Flexi DNA polymerase (C) and Invitrogen Taq DNA polymerase (D) with Bioer LifePro Thermal Cycler. The red arrow mark indicates the expected sizes of PCR amplicons.

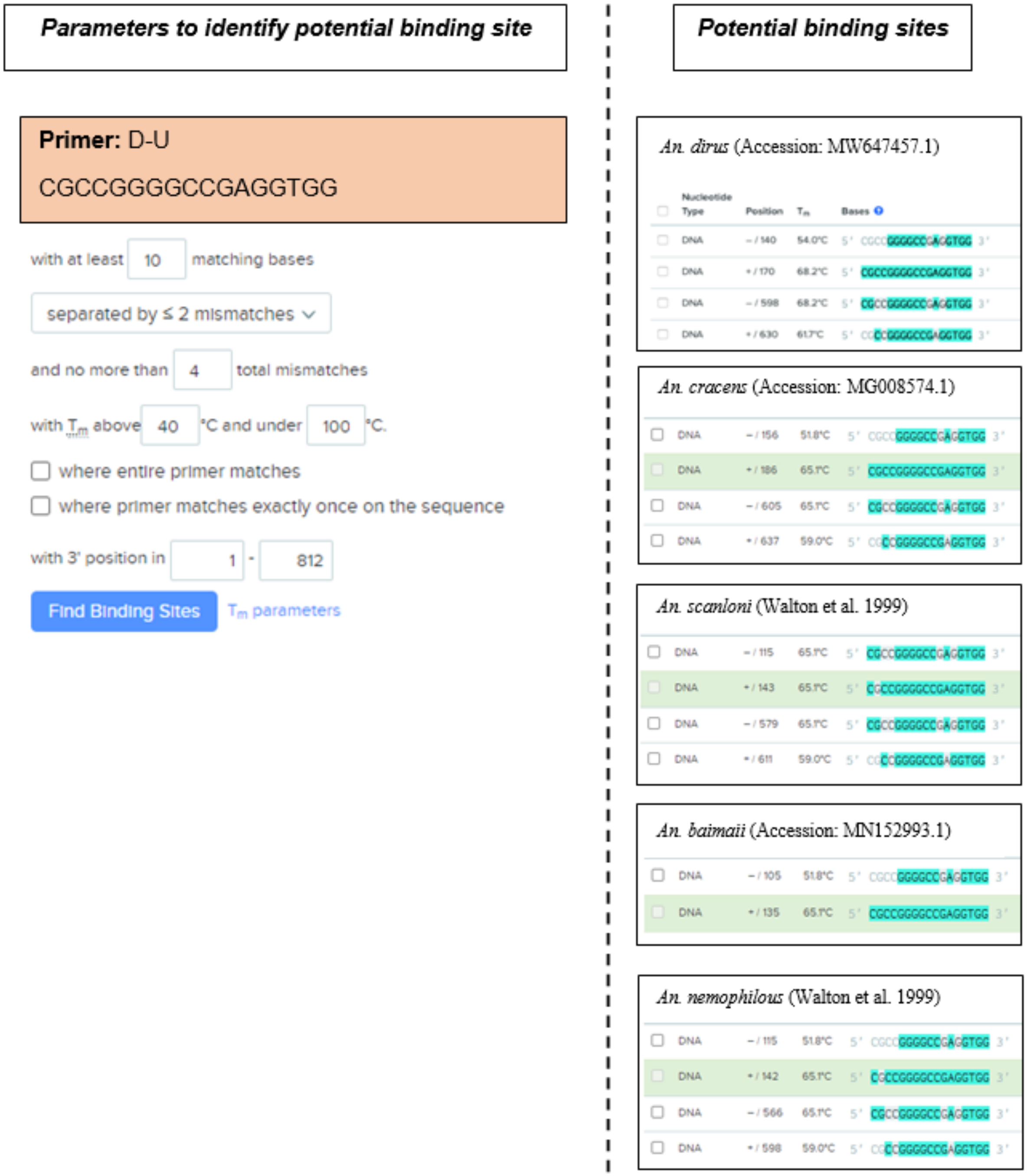


***Supplementary Figure S3. In silico* analysis reveals multiple potential binding sites of the Dirus AS-PCR D-U universal forward primer.** Potential binding sites of D-U were determined in *An. dirus*, *An. cracens, An. scanloni*, *An. baimaii*, and *An. nemophilous* ITS2 sequences using Benchling primer tool. The parameters used with this search include 1) with at least 10 matching bases, 2) separated by ≤ 2 mismatches and no more than 4 total mismatches, 3) Tm above 40°C and under 100°C. The teal color indicates a matching bases between the primer and target sites.

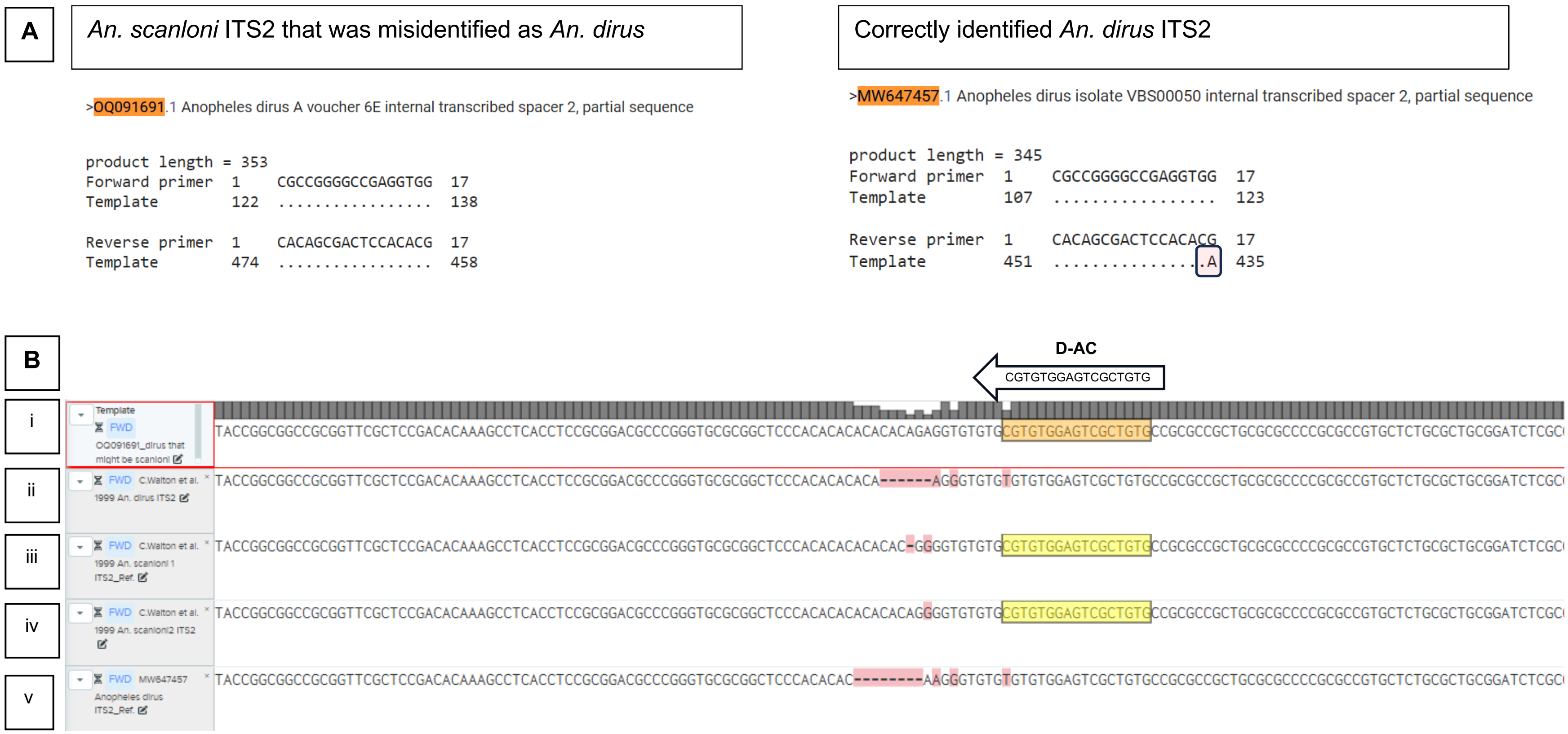


***Supplementary Figure S4. In silico* analyses reveals misidentification of *An. scanloni* as *An. dirus*.** (A) Primer-BLAST results of the D-U and D-AC primers from Dirus AS-PCR reveals potential misidentification of *An. scanloni* as *An. dirus*. Complete match of D-AC reverse primer to ITS2 sequence from the database suggests misidentification of *An. scanloni* as *An. dirus*. Single nucleotide mismatch at 3' end of D-AC reverse primer to ITS2 sequence from the database indicates that the sample is a correctly identified *An. dirus*. (B) Multiple Sequence Alignment of ITS2 sequence of *An. dirus* and *An. scanloni* demonstrated misidentification of *An. scanloni* as *An. dirus*. (i) A sequence of *An. dirus* obtained from the database which might be misidentified from *An. scanloni*. (ii) A sequence of *An. dirus* retrieved from Walton et al. (1999). (iii), (iv) A sequence of *An. scanloni* retrieved from Walton et al. (1999). (v) Sequences of *An. dirus* received from the database (MW647457). The yellow color indicates exact matches between the primer and target sites.

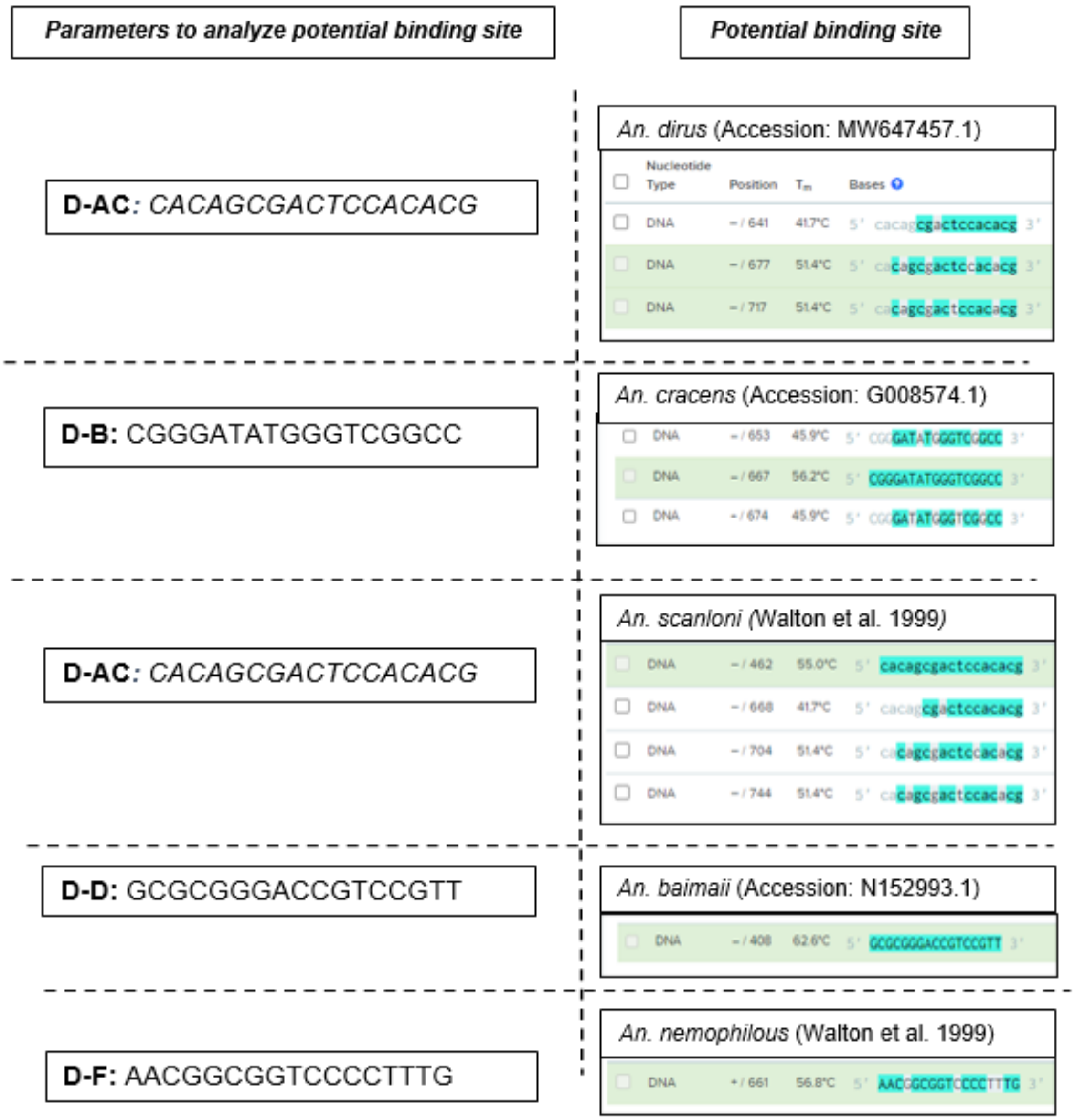


***Supplementary Figure S5***. ***In silico* analysis reveals multiple potential binding sites of the Dirus AS-PCR species specific D-AC, D-B, D-D, and D-F reverse primers.** Potential binding sites of each primer were determined in ITS2 sequences of their respective target species using Benchling primer tool. The parameters used with this search include 1) with at least 10 matching bases, 2) separated by ≤ 2 mismatches and no more than 4 total mismatches, 3) Tm above 40°C and under 100°C. The teal color indicates a matching bases between the primer and target sites

```
                    DiCSIP-Uni-Fwd
                  CTCACTACCTATGTCTCGCCC  --->
                  1                                                                              80
An. dirus         TCCTGGAT--GAGTGATGGATACAGAGCGGGCGACCGCGTCGCTGGCCTGCACGCGCACTTACGCGCCGGTCCCTGGTCG
An. cracens       TCCTGGTA--GAGTGATGGATACAGAGCGGGCGACCGCGTCGCTGGCCTGCACGCGCACTTACGCGCTGGTCCCTGGTCG
An. scanloni      TCCTGGAT--GAGTGATGGATACAGAGCGGGCGACCGCGTCGCTGGCCTGCACGCGCACTTACGCGCCGGTCCCTGGTCG
An. baimaii       TCCTGGTTGAGAGTGATGGATACAGAGCGGGCGACCGCGTCGCTGGCCTGCACGCGCACTTACGCGCTGGTCCCTGGTCG
An. nemophilous   TCCTGGAT--GAGTGATGGATACAGAGCGGGCGACCGCGCCGCTGGCCTGCACGCGCACTTACGCGCTGGTCCCTGGTCG

                  81                                                                            160
An. dirus         TTAGATGTTGCGCCCGTAGCCTTCCGAGGTCTGCAAGCGCCAGATGCTTCCGAC-ACGCACAAGGGGACCGCCGTTCGTG
An. cracens       TTAGATGTTGCGCCCGTAGCCTTCCGAGGTCTGCAAGCGCCAGATGCTTCCGACAACACACACGGGGACCGCCGTTCGTG
An. scanloni      TTAGATGTTGCGCCCGTAGCCTTCCGAGGTCTGCAAGCGCCAGATGCTTCCGAC-ACGCACAAGGGGACCGCCGTTCGTG
An. baimaii       TTAGATGTTGCGCCCGTAGCCTTCCGAGGTCTGCAAGCGCCAGATGCTTCCGAC-ACG--CAAGGGGACCGCCGTTCGTG
An. nemophilous   TTAGATGTTGCGCCCGTAGCCTTCCGAGGTCTGCAAGCGCCAGATGCTTCCGAC-ACG--AAAGGGGACCGCCGTTCGTG

                                                                                          D-D
                                                                       <---  T-TGCCTGCCAGGGCGCG
                  161
An. dirus         CGGTCGGTGCCTACAAGGTACCGGCGGCCGCGGTTCGCTCCGACACAAAGCCTCACCTCCGC-GGACGCCCGGGTGCGCG
An. cracens       CGGTCGGTGCCTACAAGGTACTGGCGGCCGCGGTTCGCTCCGACACAAAGCCCCACCTCCGCTGGACGCCCGCGGGTGCG
An. scanloni      CGGTCGGTGCCTACAAGGTACCGGCGGCCGCGGTTCGCTCCGACACAAAGCCTCACCTCCGC-GGACGCCCGGGTGCGCG
An. baimaii       CGGTCGGTGCCTACAAGGTACCGGCGGCCGCGGTTCGCTCCGACACAAAGCCTCACCTCCGA-ACGGACGGTCCCGCGCG
An. nemophilous   CGGTCGGTGCCTACAAGGTACCGGTGGCCGCGGTTCGCCTCGACACAAAGCCTCACCTCCGC-GGACGCCCGGGTGCGCG

                     DiCSIP-Fwd-C                                                     DiCSIP-Rev-F
                  CGAGGGTGTGTGTGTGTG  --->                                      <---  GCGAGACGCG
An. dirus         GCTCCCACACACA-------AGGGTGTGTGTGTGGAGTCGCTGTGCCGCGCCGCTGCGCGCCCCGCGCCGTGCTCTGCGC
An. cracens       CGACTCGTTCGT--------------------TCGAGTCGCTGTGCCGCGCCGCTGCGCGCCCCGCGCCGTGCTCTGCGC
An. scanloni      GCTCCCACACACACACACGGGGTGTGTGCGTGTGGAGTCGCTGTGCCGCGCCGCTGCGCGCCCCGCGCCGTGCTCTGCGC
An. baimaii       GGTGCTGGGCACC-------TCGTCGCGGG--------------------------------------------------
An. nemophilous   GCTCCCACACACA-------AGGGTGTGTGTGTGGAGTCGCCGTGCCGCGCCGCTGCGCGCCCCGCGCCGCGCTCTGCGC
```

48

```
                      ACGCCT
                                                                                        400
An. dirus             TGCGGA--TCTCGCGCGCTCTCGTCGCGCTCTCGCTCTCTCCCGCACCGTGGGAAGTGCAACGTTGCCACGTGGGCCCCC
An. cracens           TACGGATCTCTCGCGCGCTCTTGACGCGCTCTCGCCCTCTCCCGCACCGTGGGAAGTGCAACGTTGCCACGTGGGCCCCC
An. scanloni          TGCGGA--TCTCGCGCGCTCTCGTCGCGCTCTCGCTCTCTCCCGCACCGTGGGAAGTGCAACGTTGCCACGTGGGCCCCC
An. baimaii           --------------------------------------------------------------------------------
An. nemophilous       TGCGGA--TCTCGCGCG--CTCGTCGCGCTCTCGCTCTCTCCCGCACCGTGGGAAGTGCAACGTTGCCACGTGGGCCCCC

                                                                         D-B
                                                                 CCGGCTGGGTATAGGG--C
                      401                                                               480
An. dirus             CCGTCTGGGGCCGTGGTGGAAAGCGTTGTGCGCAGGTTGACCGACATCAGCCGCCCCATCCCGC--TC----AGCGGTGC
An. cracens           CCGTCTGGGGCCGTGGTGGAAAGCGTTGTGCGCAGGCTGACCGACATCGGCCGACCCATATCCC--GCCAAGAGCGGTGC
An. scanloni          CCGTCTGGGGCCGTGGTGGAAAGCGTTGTGCGCAGGTTGACCGACATCAGCCGCCCCATCCCGC--TC----AGCGGTGC
An. baimaii           --------------------------------------------------------------------------------
An. nemophilous       CCGTCTGGGGCCGTGGTGGAAAGCGTTGTGCGCAGGTTGACCGACATCAATCGCCCCATCCCGCAAAC----AGCGGTGC

                                                                         DiCSIP-Rev-AC
                                                                 CAACGGCCAGTCCACCTCACTA
                      481                                                               560
An. dirus             CCCGTGTGGAG-ACG--AGTGGAGAGTG----TCGCTCGAGATGCCGCGTCGAGTTGCCGGTCAGGTGGAGTGATACGTG
An. cracens           CTCGTGTGGAGAACG--GGAGGAGAGTGTCGCTCGCTCGAGCGTCGCCGACGACGATGCCGTCAGGTGGAGTG---CGCG
An. scanloni          CCCGTGTGGAG-ACG--AGTGGAGAGTG----TCGCTCGAGACGCCGCGTCGAGTTGCCGGTCAGGTGGAGTGATACGTG
An. baimaii           ---------------------------------------------------------------------------ACACA
An. nemophilous       CCCGTGTGGAG-AGAACAGGCGAGAGTG----TCGCTCGAGTGCCAACGATG------CCGTCAGGTGGAGT--------
```

***Supplementary Figure S6*. Multiple sequence alignment of ITS2 sequences of five species of the *An. dirus* complex and binding sites of primers used in DiCSIP.** The sequences of *An. dirus* (Accession: MW647457.1), *An. cracens* (Accession: MG008574.1) and *An. baimaii* (Accession: MN152993.1) were retrieved from NCBI, while those of *An. scanloni* and *An. nemophilous* were obtained from Walton et al. (1999). The correct binding sites of the primers were highlighted in red color letters.

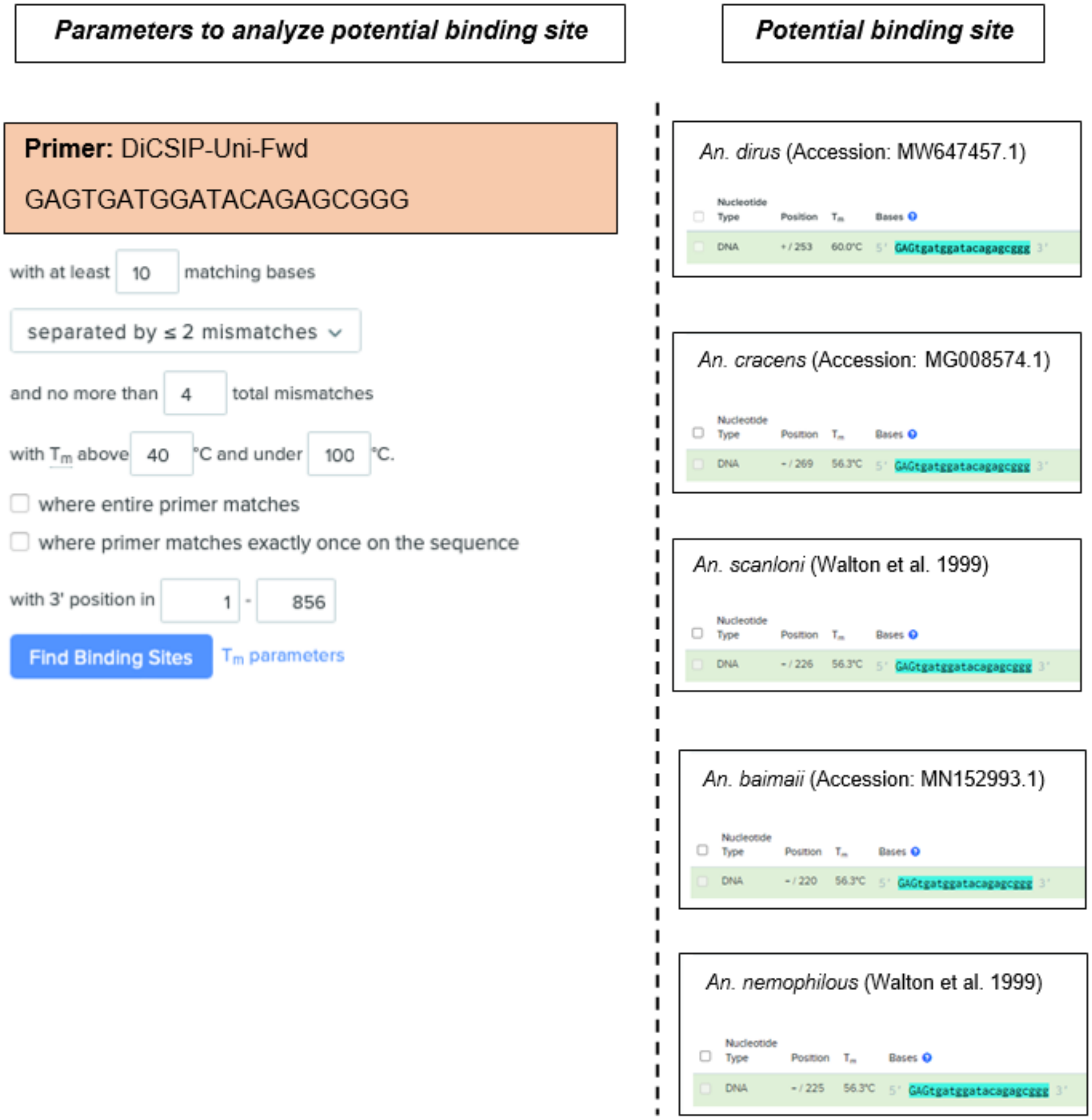


***Supplementary Figure S7*. *In silico* analysis reveals only on-target potential binding sites of the DiCSIP-Uni-Fwd universal forward primer.** Potential binding sites of DiCSIP-Uni-Fwd were determined in *An. dirus*, *An. cracens, An. scanloni*, *An. baimaii*, and *An. nemophilous* ITS2 sequences using Benchling primer tool. The parameters used with this search include 1) with at least 10 matching bases, 2) separated by ≤ 2 mismatches and no more than 4 total mismatches, 3) Tm above 40°C and under 100°C. The teal color indicates a matching bases between the primer and target sites.

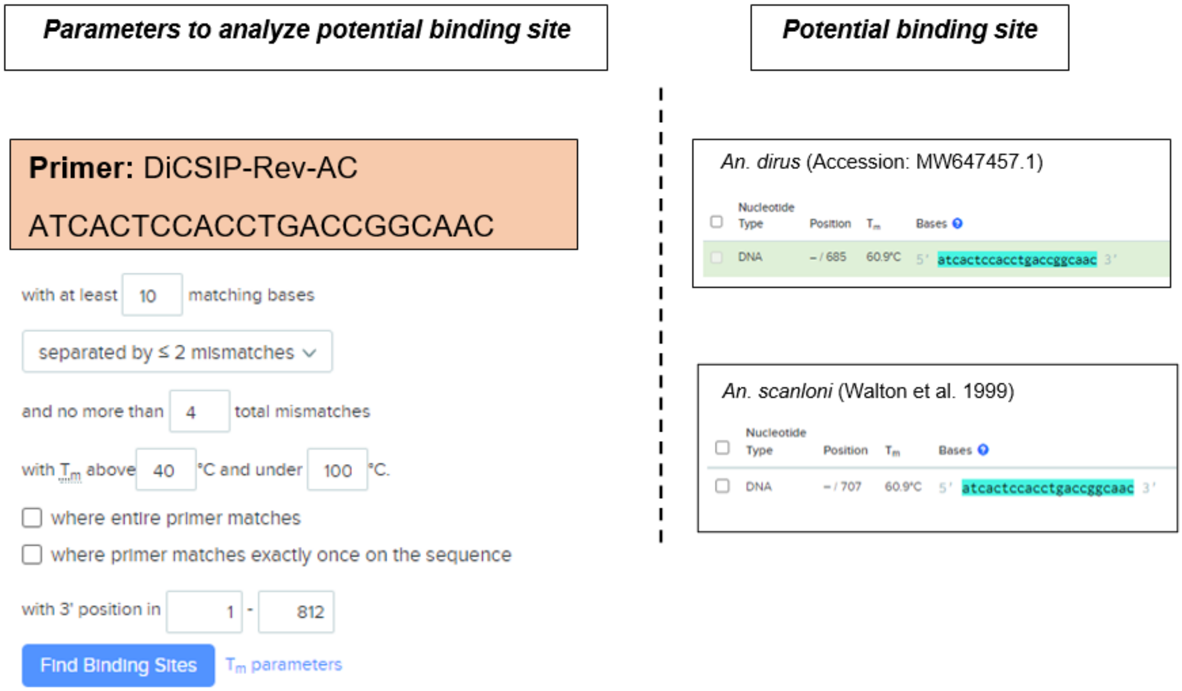


***Supplementary Figure S8. In silico*** **analysis reveals only on-target potential binding sites of the DiCSIP-Rev-AC Dirus/Scanloni specific reverse primer.** Potential binding sites of DiCSIP-Rev-AC were determined in *An. dirus,* and *An. scanloni* ITS2 sequences using Benchling primer tool. The parameters used with this search include 1) with at least 10 matching bases, 2) separated by ≤ 2 mismatches and no more than 4 total mismatches, 3) Tm above 40°C and under 100°C. The teal color indicates a matching bases between the primer and target sites.

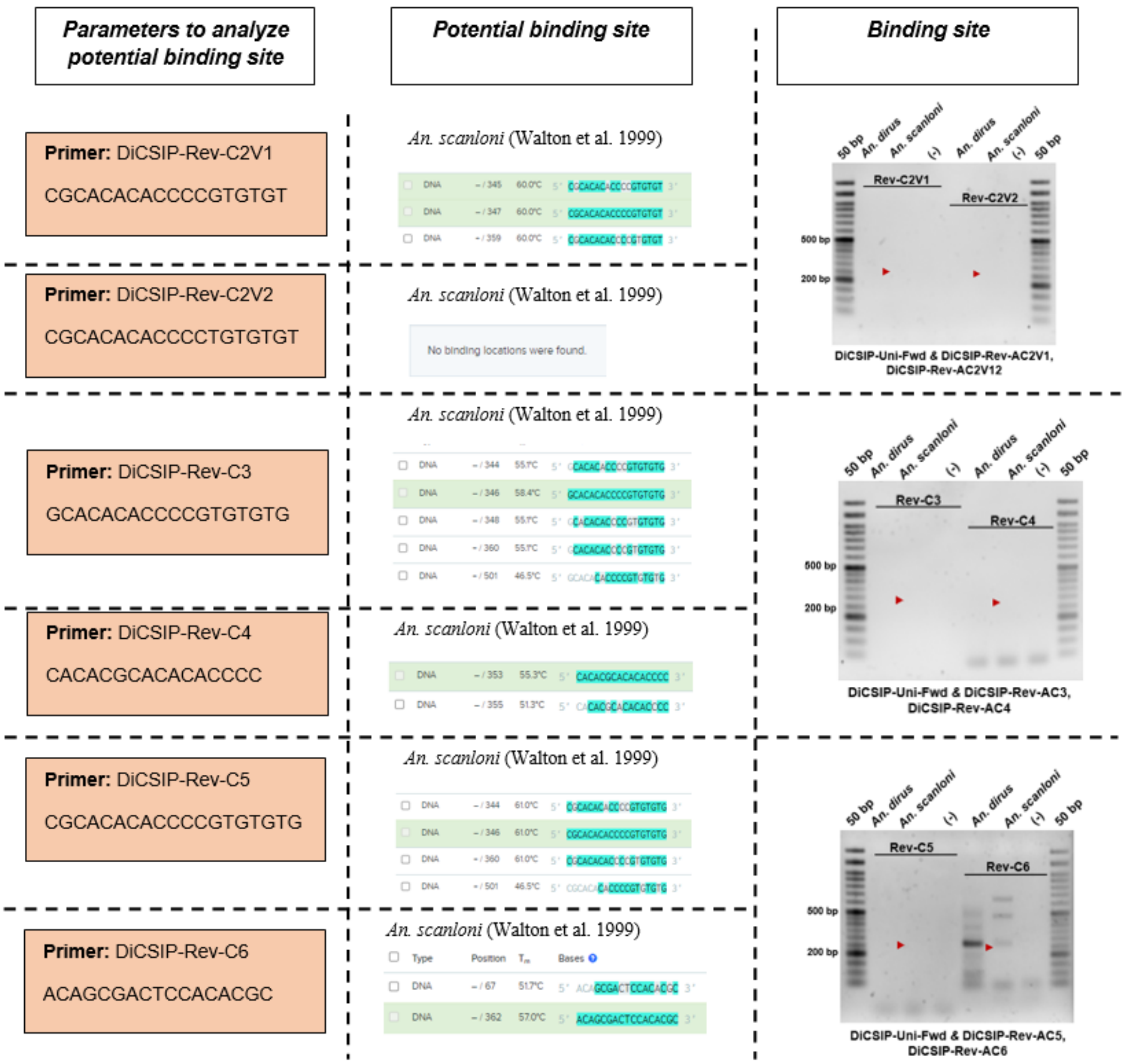


***Supplementary Figure S9*. DiCSIP-Rev-C *An. scanloni* specific reverse primers cannot be used for *An. scanloni* identification.** *In silico* analysis reveals off-target potential binding sites of the DiCSIP-Rev-C *An. scanloni* specific reverse primer. Potential binding sites of DiCSIP-Rev-C1-6 were determined in *An scanloni* ITS2 sequences using Benchling primer tool. The parameters used with this search include 1) with at least 10 matching bases, 2) separated by ≤ 2 mismatches and no more than 4 total mismatches, 3) Tm above 40°C and under 100°C. The teal color indicates a matching bases between the primer and target sites. Six new reverse primers specific to *An. scanloni* were validated using a single-plex PCR to distinguish between *An. dirus* and *An. scanloni*. The red arrow mark indicates the expected sizes of PCR amplicons.

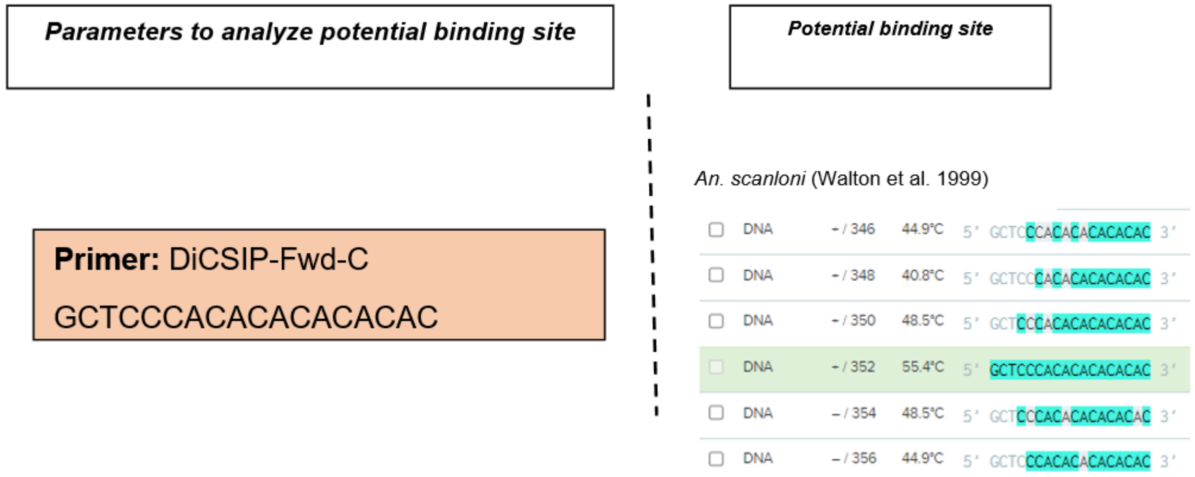


***Supplementary Figure S10.* *In silico* analysis reveals potential off-target binding sites of the DiCSIP-Fwd-C *An. scanloni* specific forward primer.** Potential binding sites of DiCSIP-Fwd-C were determined in *An. scanloni* ITS2 sequence using Benchling primer tool. The parameters used with this search include 1) with at least 10 matching bases, 2) separated by ≤ 2 mismatches and no more than 4 total mismatches, 3) Tm above 40°C and under 100°C. The teal color indicates a matching bases between the primer and target sites.

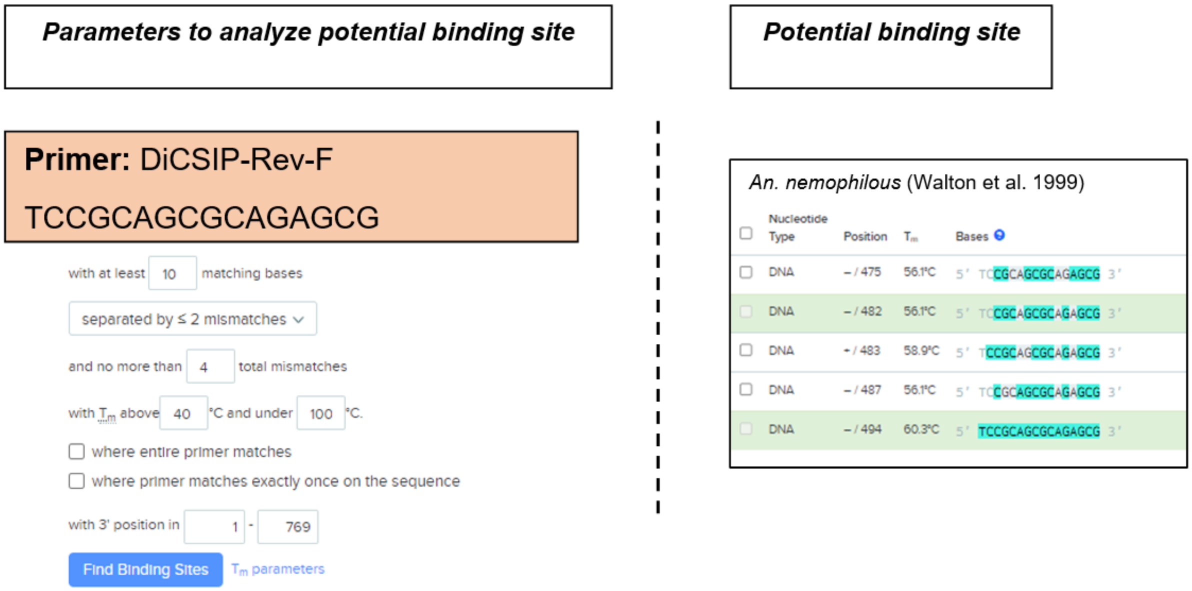


***Supplementary Figure S11.*** ***In silico*** **analysis reveals potential off-target binding sites of the DiCSIP-Rev-F** ***An. nemophilous*** **specific reverse primer.** Potential binding sites of DiCSIP-Rev-F were determined in *An. nemophilous* ITS2 sequence using Benchling primer tool. The parameters used with this search include 1) with at least 10 matching bases, 2) separated by ≤ 2 mismatches and no more than 4 total mismatches, 3) Tm above 40°C and under 100°C. The teal color indicates a matching bases between the primer and target sites.

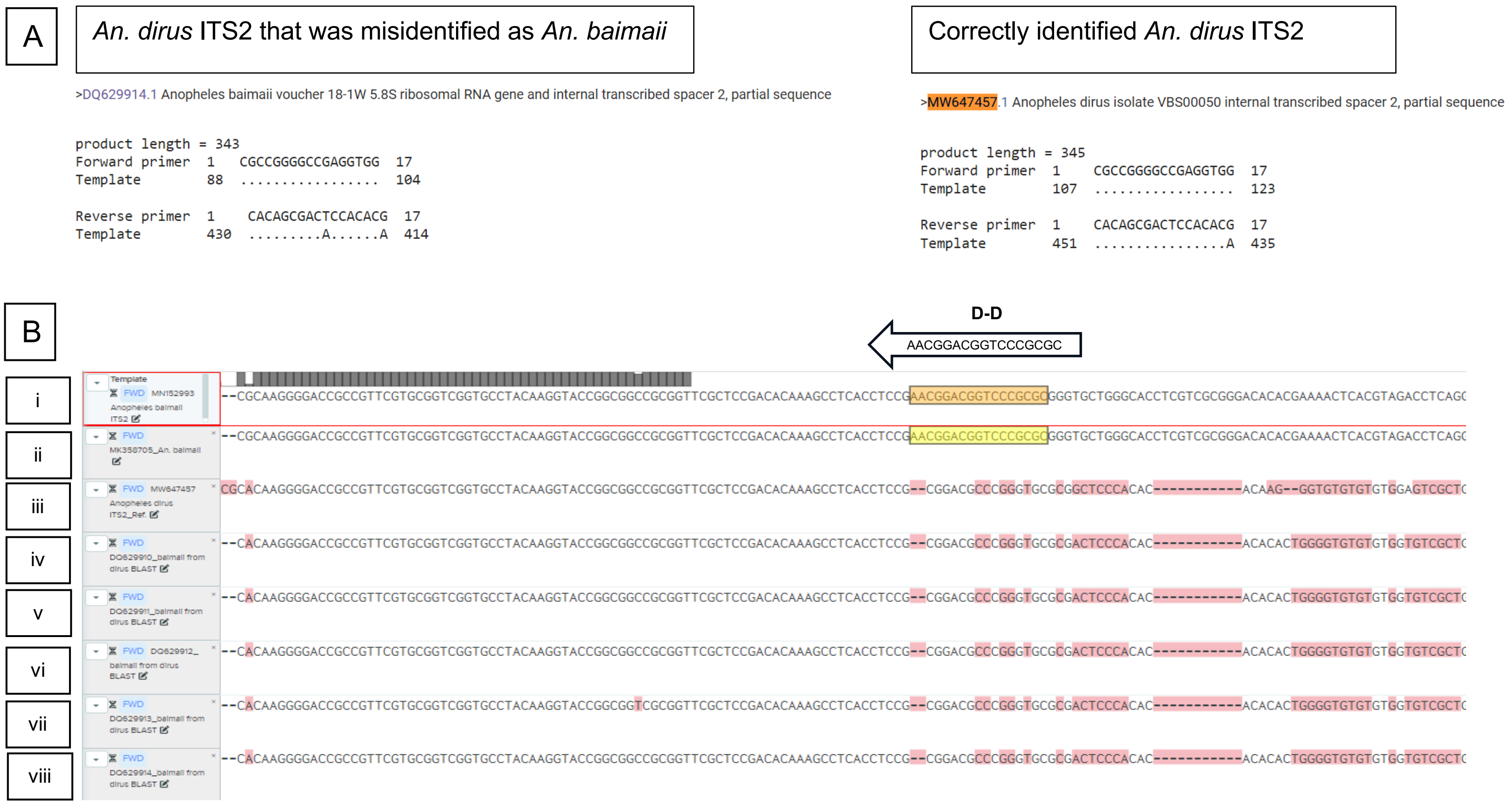


***Supplementary Figure S12.*** ***In silico*** **analyses reveal misidentification of *An. dirus* as *An. baimaii*.** (A) Primer-BLAST results of the D-U and D-AC primers from Dirus AS-PCR reveals potential misidentification of *An. dirus* as *An. baimaii*. (B) ITS2 sequence of *An. dirus* and *An. baimaii* was aligned in Benchling software with reverse D-D primer to observed potential misidentification of DNA sequence in NCBI database. (i) & (ii) Sequences of *An. baimaii* obtained from the database. (iii) A sequence of *An. dirus* retrieved from the database. (iv), (v), (vi), (vii) & (viii) Sequence of *An. baimaii* retrieved from the database which might be misidentification from *An. dirus*. The yellow color indicates a matching bases between the primer and target sites.
